# Structural and Mechanistic Insights into Symmetry Conversion in Plant GORK K^+^ Channel Regulation

**DOI:** 10.1101/2024.12.17.628833

**Authors:** Qi-yu Li, Li Qin, Ling-hui Tang, Chun-rui Zhang, Shouguang Huang, Ke Wang, Gao-hua Zhang, Ning-jie Hao, Qian Xiao, Tongxin Niu, Min Su, Rainer Hedrich, Yu-hang Chen

## Abstract

GORK is a shaker-like potassium channel in plants that contains ankyrin (ANK) repeats. In guard cells, activation of GORK causes K^+^ efflux, reducing turgor pressure and closing stomata. However, how GORK is regulated remains largely elusive. Here, we solved the cryo-EM structure of *Arabidopsis* GORK, revealing an unusual symmetry reduction (from C4 to C2) feature within its tetrameric assembly. This symmetry reduction in GORK channel is driven by ANK dimerization, which disrupts the coupling between transmembrane helices and cytoplasmic domains, thus maintaining GORK in an autoinhibited state. Electrophysiological and structural analyses further confirmed that ANK dimerization inhibits GORK, and its removal restores C4 symmetry, converting GORK to an activatable state. This dynamic switching between C2 and C4 symmetry, mediated by ANK dimerization, presents a GORK target site that guard cells regulate to switch the plant K^+^ channel between inhibited and activatable states, thus controlling stomatal movement in response to environmental stimuli.

## Main

Potassium ions are essential for plant growth and development, playing roles in osmoregulation, cell expansion, pH homeostasis, and establishing membrane potential and proton motive force^[1–2]^. In *Arabidopsis thaliana*. 71 potassium channels and transporters have been identified, among which 9 belongs to shaker-like channel family^[3]^. These channels can be categorized as outward-rectifying types, including GORK and SKOR, or inward-rectifying types, including KAT1, KAT2, AKT1, and AKT2. They sense changes in membrane potential (depolarization or hyperpolarization), regulate potassium transport, and maintain ionic homeostasis, crucial for plant growth and stress response^[2,4–7]^.

GORK, also known as <u>G</u>uard cell <u>O</u>utward-<u>R</u>ectifying <u>K</u>^+^ channel, is the only outward-rectifying potassium channel in guard cells. Since its discovery in 2000^[8]^, extensive genetic studies have demonstrated its vital role in regulating stomatal movement^[9–11]^. Stomata, formed by paired guard cells, control water loss and CO_2_ uptake in response to environmental stimuli, thereby regulating transpiration and photosynthesis^[8,12–17]^. Ion channels in guard cells control turgor pressure by mediating ion flux, which in turn regulates stomatal opening and closing^[2,5,18]^. KAT1 presents the major guard cell potassium uptake channel, required for increasing turgor pressure and volume increase promoting stomatal opening. In contrast GORK activation causes potassium efflux, reducing guard cell turgor and volume and in turn stomatal closure^[3,8–9,12,19]^.

Understanding the structure and function of these channels is critical for unraveling their roles in ion homeostasis and stomatal movement. So far, the structural study of plant KAT1 have provided important insights into its activation and role in potassium influx control^[20–21]^. However, the structure and regulatory mechanisms of GORK remain largely enigmatic. Unlike the inward-rectifying KAT1, the outward-rectifying GORK harbors a unique ankyrin (ANK) repeat domain, whose functional role in plant ion channel regulation remained yet unexplored.

In this study, we report the cryo-EM structures of *Arabidopsis* GORK (*At*GORK) in both autoinhibited state and activatable state. The full-length *At*GORK^FL^ forms a tetramer with elongated cytoplasmic regulatory domains, with its architecture showing a symmetry reduction from four-fold (C4) to two-fold (C2). Remarkable, ANK from neighboring protomers interact to form dimers, thus reducing symmetry of the tetramer and rendering GORK in an autoinhibited state. Electrophysiological studies show that mutations or truncations disrupting ANK interactions convert GORK into an activatable state. Further structural analysis of ANK-truncated *At*GORK^623^ and *At*GORK^510^ reveals that the cytoplasmic regulatory domains become highly dynamic, while strict C4 symmetry is restored in the tetrameric assembly.

Thus, our study demonstrates that ANK acts as a molecular switch for symmetry conversion (between C2 and C4), regulating GORK conformational transitions between autoinhibited and activatable states. This research provides the first comprehensive elucidation of the biological role of ANK in GORK, offering new insights into how GORK mediates stomatal closure in response to environmental signals. Furthermore, our findings broaden the understanding of ion channel regulation, particularly how symmetry changes in structural assembly regulate ion channel activity.

### Cryo-EM structure determination of the *Arabidopsis* GORK

To elucidate the structure of the GORK, we prepared *Arabidopsis* GORK (fused with an N-terminal GFP tag) from HEK293 cells and conducted single particle cryo-EM analysis. After solubilization in 1.0% n-Dodecyl-B-D-Maltoside (DDM) and 0.02% cholesteryl hemisuccinate (CHS), the *At*GORK proteins underwent purification through strep-tactin affinity chromatography, GFP removal via TEV enzyme cleavage, and subsequent gel-filtration chromatography. The final solution contained 0.006% detergent GDN, 150 mM NaCl, and 20 mM HEPES-Na pH 6.5. The most monodisperse peak fractions were pooled and concentrated to approximately 3.5 mg/mL for cryo-EM grid preparation.

We determined the cryo-EM structures of the full-length *At*GORK (*At*GORK^FL^, residues 1 to 820) in an auto-inhibited state, along with two ANK-truncated versions (*At*GORK^623^, residues 1 to 623; *At*GORK^510^, residues 1 to 510) (Table S1, Supporting Information). Tetrameric reconstructions of *At*GORK were obtained in two conformational states at resolutions of 3.4 Å (*At*GORK^FL1^) and 4.3 Å (*At*GORK^FL2^), allowing *de novo* modeling of 662 of 820 amino acids (residues 58 to 720) for each protomer, while the remaining regions were not resolved due to their intrinsic flexibility (Figure S1, Supporting Information). For the truncated *At*GORK^510^ and *At*GORK^623^, cryo-EM maps at 3.4 Å and 3.2 Å resolution, respectively, were obtained. Both maps showed high-quality density in their transmembrane domain, allowing accurate modeling, while the flexible cytosolic domains remained unmodeled due to their invisibility (Figure S2, Supporting Information). Structure-based sequence alignment for 24 representative GORKs from both monocots and dicots is shown in Figure S3, Supporting Information.

### Architecture of *Arabidopsis* GORK in symmetry-reduced conformation

*Arabidopsis* GORK (*At*GORK) forms a homo-tetrameric structure around a central pore, with a dimension of 110 Å (length) × 110 Å (width) × 170 Å (height) (Figure 1A-C). Each protomer has an N-terminal transmembrane domain (TMD, residues 58-313, containing helices S1-S6), followed by three cytoplasmic domains: the C-linker (residues 314-365), CNBDH (residues 366-510), and ANK (residues 511-724). The segment preceding the TMD (residues 1-57) and the last portion (residues 725-824, containing a putative KHA domain enriched in hydrophobic and acidic residues) are not visible in the structure, likely due to their intrinsic flexibility or the flexibility of the connecting regions.

**Figure 1.**
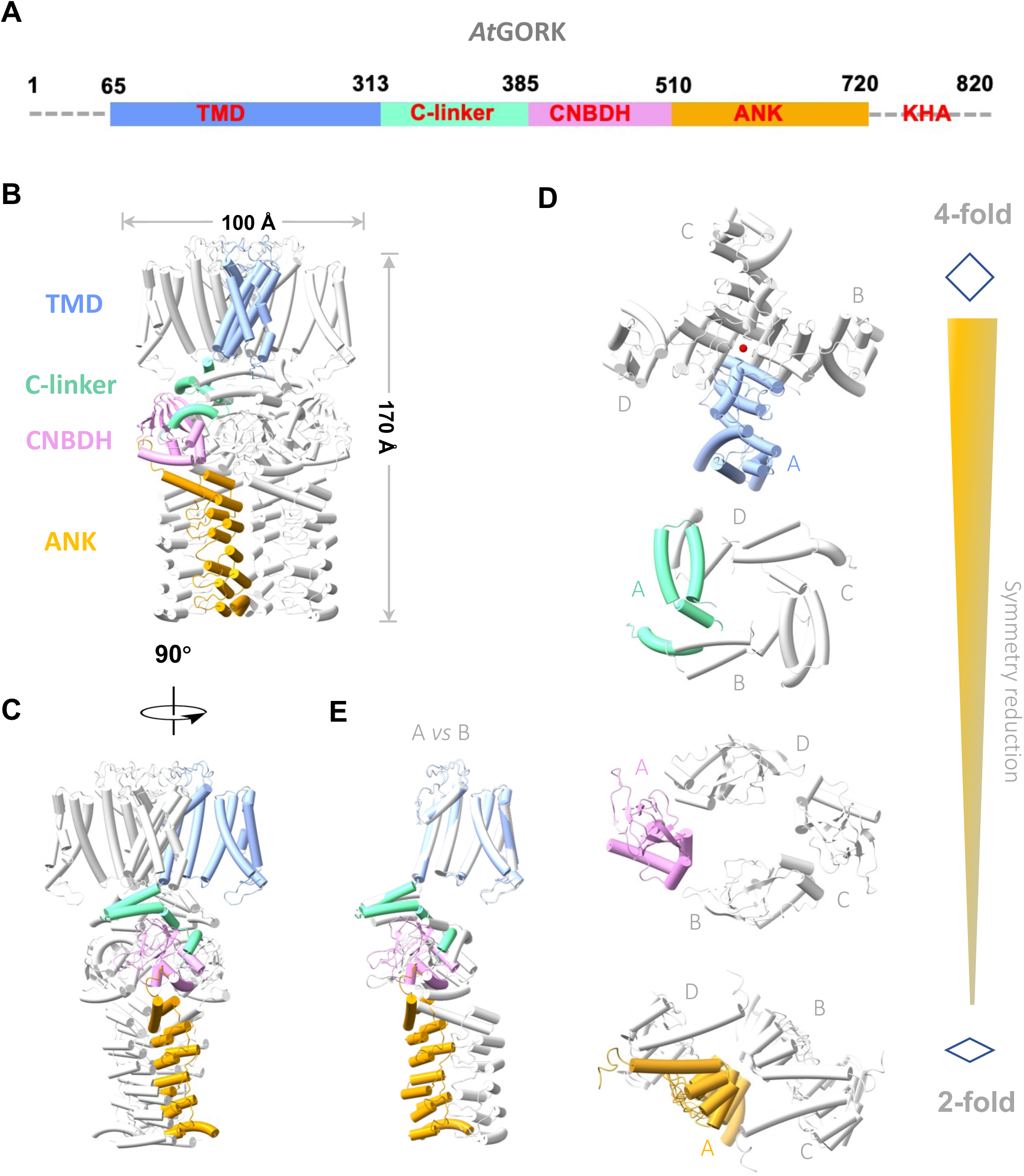
Architecture of the Arabidopsis GORK. A) Schematic diagram of the domain architecture in Arabidopsis GORK (*At*GORK). The transmembrane domain (TMD, blue), the C-linker domain (C-linker, Cyan), the cyclic nucleotide-binding domain homolog (CNBDH, purple) and the Ankyrin domain (ANK, orange) are shown. B, C) The cartoon drawing of the *At*GORK tetramer. One protomer is colored as indicated in (A), and other protomers are colored in grey. D) Symmetry reduction (from C4 to C2) within the tetrameric assembly. This represents a unique structural feature of the GORK family and likely serves as a unique mechanism for channel regulation. The current *At*GORK structure likely represents an autoinhibited, pre-activation state. Different protomers are indicated with letter A-D. E) Structural superimposition of protomer A with its neighboring protomer B.

The *At*GORK structure, unlike other tetrameric ion channels such as KAT1, exhibits a unique architectural assembly^[20]^. Notably, the symmetry in the TMD portion adopts a typical four-fold (C4), but reduces to a two-fold (C2) in their following cytoplasmic domains, thus leading to a deformation within the tetrameric channel (Figure 1D). Superimposition of its protomer reveals distinct conformations, with notable rotations and displacements occurred in the C-linker, CNBDH, and ANK domains (Figure 1E). This symmetry reduction within the tetrameric assembly appears to arise from ANK dimerization (will discuss later). To the best of our knowledge, this symmetry reduction in channel assembly is exceptionally rare and leads to deformation of the tetrameric channel. This represents a unique structural feature of the GORK family and likely serves as a unique mechanism for channel regulation. Thus, the current *At*GORK structure likely represents an autoinhibited, pre-activation state.

The TMD consists of a voltage sensing domain (VSD, helices S1–S4) and a pore-forming domain (PD, helices S5 and S6) (Figure 2A). The VSD is connected to the PD through a short linker, resulting in a “non-domain-swapped” configuration where the VSD and PD of the same subunit form a cohesive bundle. This differs from the canonical domain-swapped architecture in Shaker-like^[22]^ and Na_V_ channels^[23]^, where the VSD and PD are separated by a long S4−S5 linker. The P-loop segment of *At*GORK contains a highly conserved motif (^271^TVGYG^275^), similar to that of bacterial KcsA^[24]^, mammalian HCN^[25]^, and plant outward-rectifying SKOR^[26]^ and inward-rectifying KAT1^[20]^ and AKT1^[27]^. Structural comparisons reveal that they form a similar K^+^ selectivity filter (Figure S4A, Supporting Information). Analysis using the HOLE^[28]^ suggests that the *At*GORK adopts a closed conformation. Below the selectivity filter lies the pore gate region consists of residues on the S6 helix, with I303 and T307 creating narrowest constrictions (radius of approximately 1 Å). The S4 helix contains conserved positively charged residues (R174, R177, R179, and R187), and adopt a resting “up” conformation, similar to that in plant inward-rectifying KAT1^[20]^ and AKT1^[27]^ (Figure S4B, Supporting Information).

**Figure 2.**
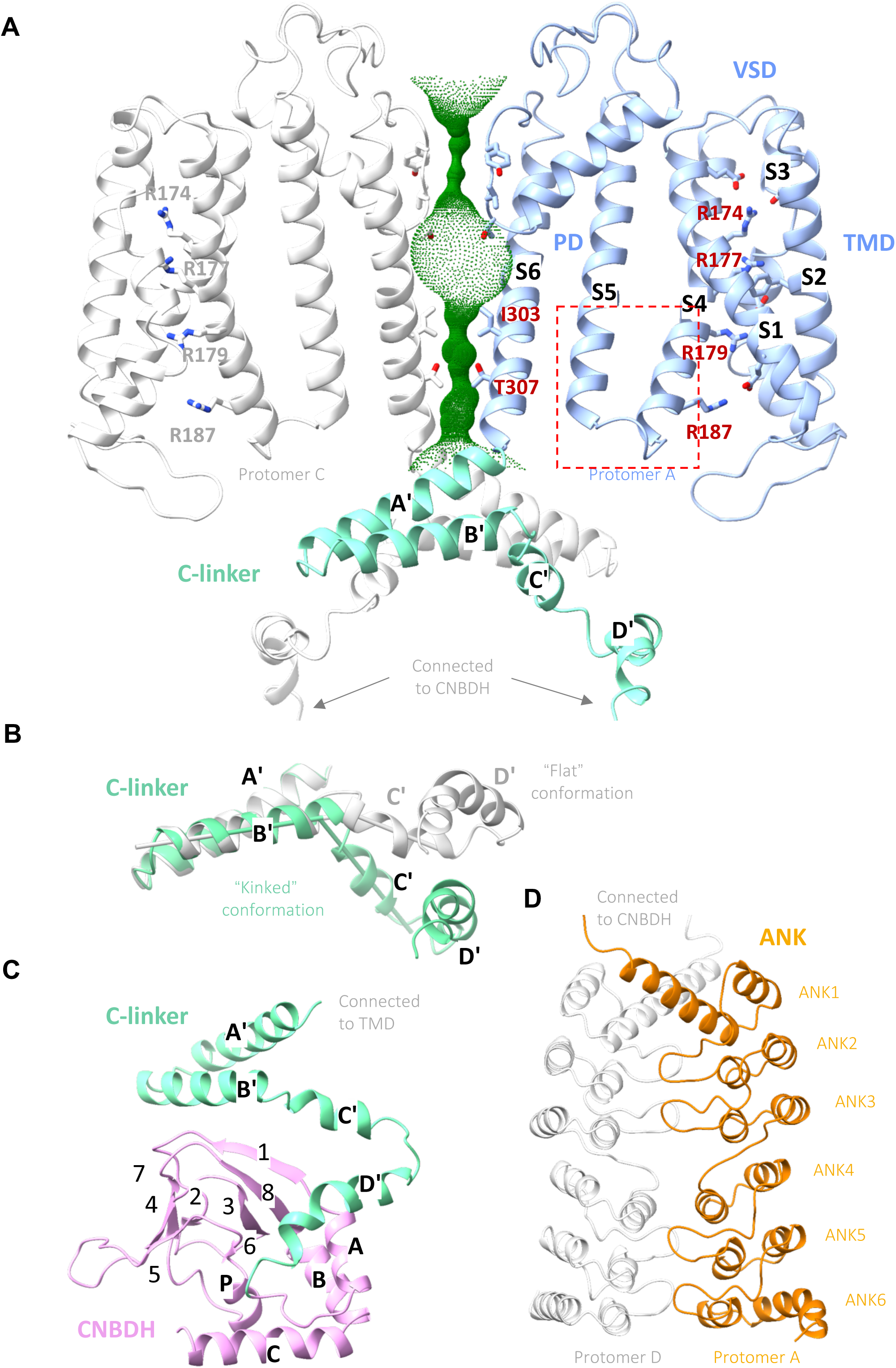
Structural domains of *At*GORK: TMD, C-linker, CNBDH and ANK. A) Ribbons drawing of the TMD/C-linker portion, with two protomers (A and C) are shown, colored and oriented as in Figure 1C. The VSD is connected to the PD through a short linker, resulting in a “non-domain-swapped” configuration. The pore-lining surface was computed by the program HOLE and drawn into a cartoon model of the GORK channel pore in a closed conformation. Critical residue in VSD (R174, R177, R179, and R187), selectivity filter (^271^TVGYG^275^) and the narrowest constrictions (I303 and T307) are shown in sticks. The S4 helix adopt a resting “up” conformation, similar to that in plant inward-rectifying KAT1 and AKT1. B) Structural comparison of the C-linker of protomer A (“kinked” conformation) with protomer B (“flat” conformation). The “flat” conformation of the C-linker actually disrupts the formation of a C4 symmetrical gating ring, resulting in decoupling between the transmembrane helices with the cytoplasmic domains, ultimately affecting channel gating. C) Ribbons drawing of the C-linker/CNBDH portion. D) Ribbons drawing of the dimeric ANK portion. ANKs from two neighboring protomers tightly interact, creating a buried area of ∼1500 Å^2^. The first three and the last three repeats are tandem units. Repeats 1-2/4-5 are canonical α-helix/α-helix/β-hairpin motifs, while repeat 3/6 lacks the β-hairpin. The presence of the ANK domain and its dimerization are unique features of the GORK family. ANK dimerization leads to symmetry reduction in the tetrameric channel. This, in turn, affects the coupling between the transmembrane helices and cytoplasmic domains, ultimately holding the ion channel in an autoinhibited, pre-activation state.

After the TMD, the C-linker is positioned next to the helix bundle crossing, connecting the transmembrane helices to the cytoplasmic domains. It comprises four helices (A’-D’), forming two “helix-turn-helix” motifs. These helices further assemble into a gating ring architecture within the tetrameric channel through “elbow-on-shoulder” inter-subunit interactions, where helix A’-turn-helix B’ (elbow) of one protomer interacts with helix C’-turn-helix D’ (shoulder) of its neighboring protomer^[29]^ (Figure 1C). Due to the reduced symmetry within the tetramer, the C-linkers exhibits distinct conformations between adjacent protomers, either “flat” or “kinked” (Figure 2B). Consequently, the C-linker in *At*GORK deviates from C4 symmetry, and the presence of the “flat” conformation disrupts the formation of a C4 symmetrical gating ring, leading to decoupling between the transmembrane helices and the cytoplasmic domains, ultimately affecting channel gating.

The CNBDH domain, connected to the C-linker, shares approximately 20% sequence identity with canonical cyclic nucleotide-binding domains (CNBDs) (Figure S5A, Supporting Information), which play crucial roles in regulating channels, kinases and transcription factors upon binding to secondary messengers (cAMP or cGMP). Despite low sequence homology, the CNBDH share a similar structural fold to those of the CNBDs (Figure S5B, Supporting Information). The CNBDH of GORK consists of four short α-helices (A, P, B, and C) and an eight-stranded β-roll (β1-β8) (Figure 2C). Helix A is located before the β-roll, Helix P is inserted within the β-roll (between β6 and β7), and Helices B and C are positioned afterward. Helices A and B interact in an antiparallel manner, and the putative phosphate-binding cassette comprises β6, Helix P, and β7.

Although the CNBDH shares overall structural similarity with animal CNBDs, there are notable differences (Figure S5B, Supporting Information). GORK lacks conserved residues for cAMP binding, such as R561 in 7lfx or R620 in 2ptm, which is replaced by Q465. More strikingly, alanine at the binding pocket entrance (A563 in 7lfx^[30]^, or A622 in 2ptm^[29]^) is replaced by bulky phenylalanine (F467 in *At*GORK), likely blocking nucleotide access. Helix C also adopts a closed conformation, with its adjacent loop obstructing the binding site entrance, suggesting that large conformational changes would be required for cyclic nucleotide binding.

Following the CNBDH, six ankyrin repeats fold into a single ANK domain, forming a slightly curved solenoid structure (Figure 2D). The first three repeats (ANK1-3) and the last three repeats (ANK4-6) are tandem units with ∼34% sequence identity. In each tandem unit, two repeats are canonical, consisting of a pair of anti-parallel α-helices and an intervening “finger” loop, while the third is a degenerate repeat lacking the “finger” loop. Remarkably, ANK repeats from two neighboring protomers forms a stable dimer through specific interactions mediated by their protruding loops, creating a buried area of ∼1500 Å^2^. This dimeric interaction is specific, corroborated by another structure of the full-length of *At*GORK^FL2^, which retains similar dimeric units despite of variation in their tetrameric conformations (Figure S1E, Supporting Information). A similar ANK dimerization pattern has also been observed in SKOR (Figure S1F, Supporting Information)^[26]^, suggesting that this dimerization is a common structural feature in outward rectifying plant ANK-containing K^+^ channels.

As a result of ANK dimerization, positional deviations and conformational changes are induced in the adjacent CNBDH and C-linker, leading to a reduction in symmetry within the tetrameric channel. This reduction affects the communication and coupling between cytoplasmic domains and transmembrane helices, ultimately maintaining GORK in an autoinhibited, pre-activation state. The structures of GORK and SKOR with reduced symmetry provide crucial insights into how the ANK domain regulates these channels, maintaining them in an autoinhibited conformation under resting conditions. This is supported by the observation of basal activity of full-length *At*GORK.

### ANK dimerization maintains GORK channel in an auto-inhibited state

The ANK repeats and their dimeric interactions are distinctive features of the GORK channel, playing a key role in reducing symmetry within its tetrameric assembly. To explore their function, we conducted trunctional analysis of the ankyrin repeats in *At*GORK expressed in *Xenopus oocytes* and inspected via TEVC electrophysiology (Figure 3A). Full-length *At*GORK produced basal currents, indicating that a small fraction of these channels was in an activatable state. Removing the KHA domain while retaining the intact ANK domain in *At*GORK (Δ737-820, residues 1-736) led to a slight increase in oocytes currents, whereas deleting the last three C-terminal ANK repeats (Δ624-820, residues 1-623) significantly enhanced current amplitude—reaching 13.3-fold that of the full-length channel at +70 mV (Figure 3B, C). However, removing all six ANK repeats produced more complex outcomes depending on the truncation site: Δ544-820 (residues 1-543) further increased current amplitude to approximately 26.3-fold that of the wild-type full-length channel, while Δ528-820 (residues 1-527) decreased current amplitude, and Δ511-820 (residues 1-510) nearly abolished the currents (Figure 3B, C).

**Figure 3.**
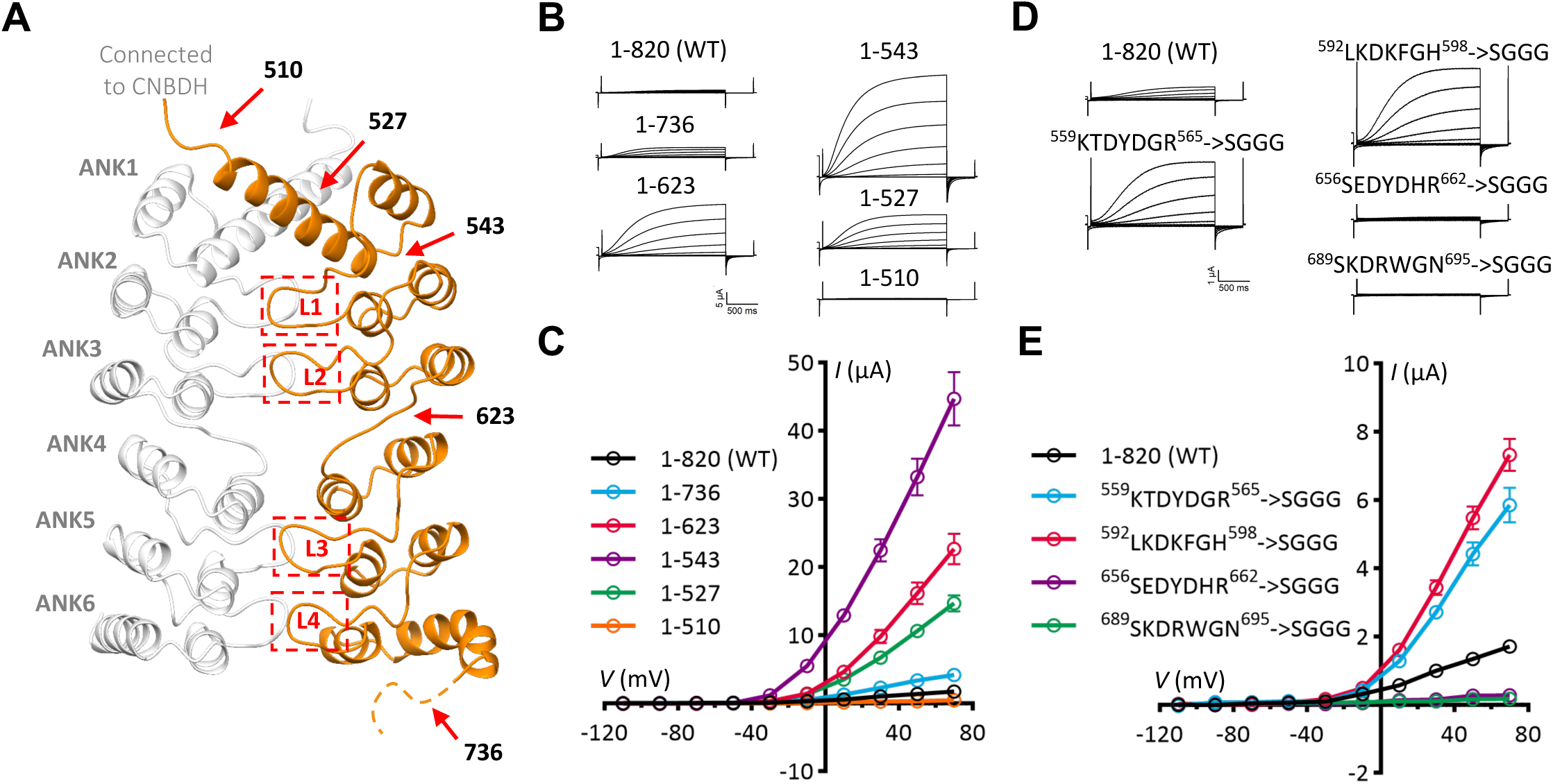
ANK plays an inhibitory role in regulating *At*GORK. A) Ribbons drawing of the ANK dimeric domain, with truncated points or mutated loops are highlighted with red arrows or in red boxes. B, C) The C-terminal truncation of *At*GORK, with representative current traces in (B) and steady-state current-voltage (*I-V*) relations in (C). Truncating half ANK domain (Δ624-820, residues 1-623) or the entire ANK domain (Δ544-820, residues 1-543) significantly increased currents. However, a further truncation (Δ511-820, residues 1-510) disrupted the channel activity. D, E) The loop-substitution in ANK of *At*GORK, with representative current traces in (D) and steady-state current-voltage (*I-V*) relations in (E). Substitution of loops between repeats 1/2 and 2/3 with “SGGG” significantly enhanced GORK activity. However, similar replacement of the loops between repeats 4/5 and 5/6 impaired GORK function. Data are mean ± SEM, n ≥ 8.

To rule out the possibility that these changes in current were due to trafficking defects, we fused a GFP tag to the N terminus of these ANK truncation mutants and observed that their surface expression levels were comparable to that of the wild-type channel (Figure S6A, B, Supporting Information). Therefore, the enhanced currents in the truncation mutants are primarily due to the disruption of ANK dimerization, which in turn releases *At*GORK from its autoinhibited state. These findings indicate that ANK dimerization is responsible for maintaining the autoinhibited state of *At*GORK, and the channel becomes activatable upon autoinhibition release. Moreover, the helix from the first ANK repeat (residues 520–543), which is directly connected to the CNBDH, is essential for the transition of *At*GORK from the autoinhibited to the activated state.

The dimerization of ANK is driven by hydrophobic interactions mediated by the hairpin loops protruding from the ankyrin repeats. Inspired by this structural insight, we replaced the hairpin loop with an unrelated short “SGGG” motif. Notably, substituting the loops between repeats 1/2 and 2/3 significantly enhanced *At*GORK activity, whereas similar replacement between repeats 4/5 and 5/6 impaired *At*GORK function (Figure 3D, E). To rule out the possibility that these mutations affected channel trafficking, we also performed GFP fusion analysis on the resutant mutants (Figure S6A, B, Supporting Information), which confirmed normal surface expression levels. This aligns well with findings from ANK truncational studies (Figure 3B, C), confirming that disrupting ANK dimeric interactions release *At*GORK from autoinhibition and thus allowing channel activation. However, these observations also suggest that the ANK repeats in GORK have a more complex role than initially anticipated: they are not only essential for maintaining the autoinhibited state but also play a pivotal role in stabilizing the open state during channel activation.

### Mutational test on conserved acidic residues at the C-linker/TMD interface

The C-linkers assemble into a disc-shaped gating ring via “elbow-on-shoulder” interactions^[29]^, directly contacting the TMD and playing a crucial role in channel gating. Conserved negatively charged residues, including E317, D321, and D325, are positioned on the C-linker A’ helix, forming a negatively charged ring at the entrance of the ion-conducting pathway (Figure 4A) and engaging in an intricate interaction nexus with conserved K/R from the TMD (Figure 4B). Specifically, E317 and D321 (C-linker of protomer A) form ionic interactions with K312 (Helix S6’ of protomer D), while R200 (Helix S5 of protomer A) forms a salt bridge with E317 from its adjacent protomer B. In such a fashion, the C-linker-mediated gating ring is connected with the VSD and PD within the TMD through their domain-swapped interfaces. These three residues correspond to conserved R, D, and A in the inward K^+^ channels KAT1^[20]^ and AKT1^[27]^, respectively.

**Figure 4.**
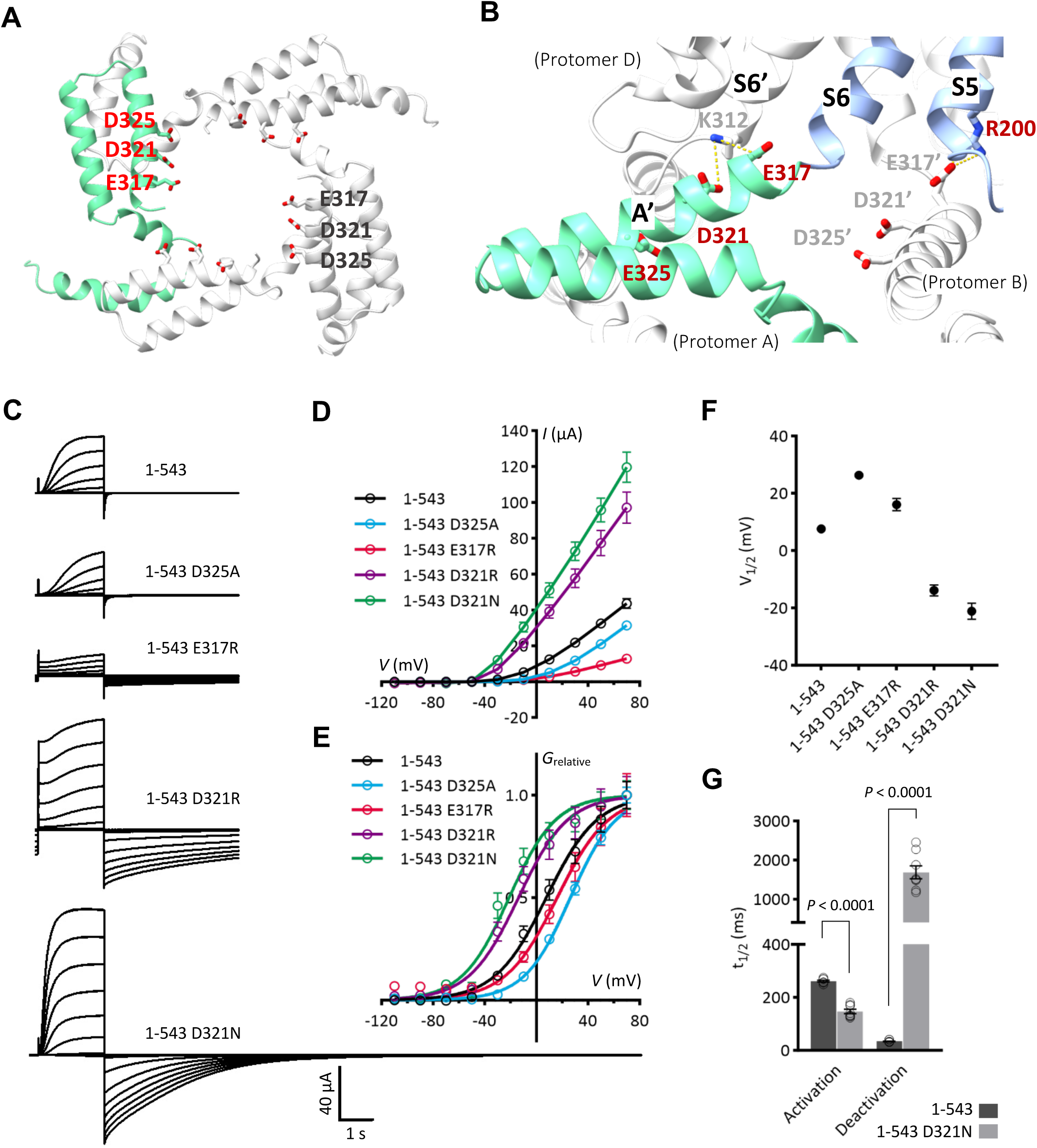
Mutational tests on conserved acidic residues at the C-linker/TMD interface. A) Ribbons drawing of the C-linkers in tetrameric assembly, highlighting negatively charged residues (E317, D321, D325) in sticks. B) Close-up view of the interface between the C-linker and TMD. E317 and D321 (C-linker of protomer A) form ionic interactions with K312 (Helix S6’ of protomer D), while R200 (Helix S5 of protomer A) forms a salt bridge with E317 from its adjacent protomer B. C-G) Electrophysiological analyses of E317, D321 and D325 mutations in *At*GORK^543^. Representative current traces (C) and steady-state current-voltage (*I-V*) relations (D) are shown. Relative conductance-voltage (*G_relative_-V*) curves (E) and half-activation voltage (V_1/2_) values (F) were generated through Boltzmann sigmoidal fitting (outliers excluded). Conductance was calculated using the equation *G* = *I*/(*V - E_K_*), where *I* is the steady-state current, *V* is the test potential, and *E_K_* (–58 mV) was derived from the Nernst equation based on intracellular (∼100 mM) and extracellular (10 mM) K^+^ concentration in the oocyte TEVC recordings. Relative conductance was calculated by normalization to the maximal conductance. Half-time (t_1/2_) for activation at +70 mV and deactivation at −110 mV, calculated as ln (2)·τ where τ is the time constant from single-exponential decay fitting, is shown in (G). Data are mean ± SEM, n ≥ 8. Significance analysis was performed using unpaired Student’s t-test, with P-values displayed on the bar charts.

To further validate their functional role, we introduced the mutations, E317R, D321 (N or R), and D325A into both full-length *At*GORK and the ANK-truncated *At*GORK^543^ construct. In full-length GORK, E317R and D325A enhanced current amplitude, whereas D321N and D321R produced an even larger increase (Figure S7A, B, Supporting Information). In contrast, in the activatable ANK-truncated *At*GORK^543^, E317R and D325A partially reduced the current amplitude, yet it remained higher than that of the wild-type full-length *At*GORK, while D321N and D321R still significantly enhanced the currents (Figure 4C, D). In addition, these mutants exhibited distinct alterations of half-activation voltage (V_1/2_) in their *G_relative_–V* relationships: V_1/2_ of E317R negatively shifted in full-length *At*GORK but showed slightly positively shifted in *At*GORK^543^; V_1/2_ of D325A had no apparent shift in the full-length construct but positively shifted in GORK^543^; and V_1/2_ of D321N and D321R negatively shifted in both constructs (Figure 4E, F; Figure S7C, D, Supporting Information).

Intriguingly, E317R and both D321 mutations (D321N and D321R) exhibited significant changes in deactivation kinetics. Specifically, when the membrane potential was held at −110 mV following the test voltage, the tail current decay was markedly delayed (Figure 4C; Figure S7A, Supporting Information). For D321N, the deactivation time constant (t_1/2_) was prolonged in both the full-length GORK and the ANK-truncated *At*GORK^543^, increasing from 75 ms to 1992 ms, and from 34 ms to 1687 ms, respectively (Figure 4G; Figure S7E, Supporting Information). Moreover, D321N accelerated the activation kinetics, with t_1/2_ decreasing from 462 ms to 286 ms in the full-length, and from 261 ms to 147 ms in the truncated form respectively (Figure 4G; Figure S7E, Supporting Information). Notably, inward K⁺ tail currents were observed during the −110 mV holding phase, indicating that these mutations disrupted the original outward rectification, thereby allowing inward K⁺ flow.

### Disrupting ANK-dimerization relieves constraints and restores C4 symmetry

Our structural studies reveal that ANK dimerization in full-length *At*GORK is directly responsible for reducing its tetrameric symmetry from C4 to C2, maintaining an autoinhibited state. Under resting conditions, GORK shows minimal basal currents, whereas ANK-truncated mutants, including *At*GORK^623^ and *At*GORK^543^, show significantly increased currents, suggesting that ANK truncations relieve autoinhibition and enhances GORK activity. Structural analysis of *At*GORK^623^ and *At*GORK^510^ only resolved the TMD portion, with their cytoplasmic regions being too flexible for visualization (Figure S2, Supporting Information). These findings suggest that ANK truncation relieves the autoinhibitory constraints imposed by ANK dimerization, converting *At*GORK into an activatable state.

Structural analysis also provided some clues regarding the restoration of C4 symmetry in the *At*GORK. Firstly, the structures of *At*GORK^623^ and *At*GORK^510^ are highly similar, with a root-mean-square deviation (RMSD) of approximately 0.48 Å for their superimposed tetramers (Figure 5A). Importantly, the ANK-truncated protomers exhibit strict C4 symmetry, with RMSDs of approximately 0.16 Å/256 superimposed Cα. In contrast, the TMD of full-length *At*GORK^FL1^ displays quasi-C4 symmetry (essentially C2 symmetry), with subtle yet distinct differences between protomers A/C and B/D at the N-terminal end of helix S5, yielding an RMSD of approximately 0.75 Å/256 superimposed Cα (Figure 5B, C). A comparison of protomers A and B in both *At*GORK^FL1^ and *At*GORK^623^ reveals that the protomers in *At*GORK^623^ closely resembles protomer A in *At*GORK^FL1^, rather than protomer B (Figure 5D). This suggests that ANK dimerization results in C2 symmetry in the full-length *At*GORK^FL1^, whereas ANK truncation restores C4 symmetry in the *At*GORK^623^.

**Figure 5.**
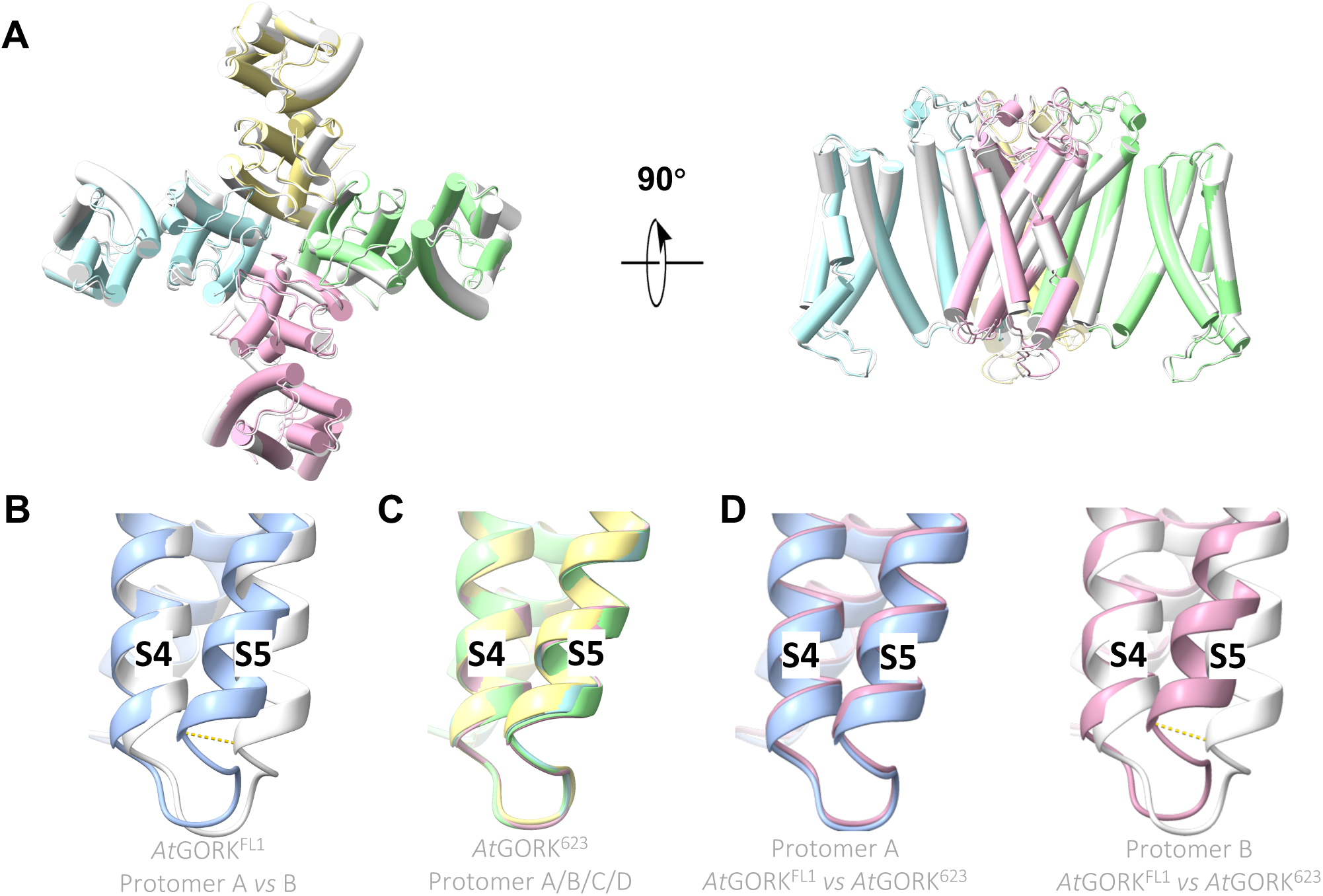
Structural comparison of the auto-inhibited *At*GORK and the ANK-truncated variants. A) Superimposition of the ANK-truncated variants (*At*GORK^623^ and *At*GORK^510^), with each protomer of *At*GORK^623^ shown in different color, while all four protomers in *At*GORK^510^ are displayed in grey. Both truncated mutants showed nearly identical structures, with a root-mean-square deviation (RMSD) of approximately 0.48 Å for their superimposed tetramers. B-D) Close-up views of S4-S5 helix regions in the superimposed protomers of *At*GORK^FL1^ and *At*GORK^623^. (B) compares protomer A (blue) and protomer B (grey) of *At*GORK^FL1^, while (C) shows all four protomers of *At*GORK^623^. These comparisons reveal that *At*GORK^FL1^ exhibits quasi-C4 symmetry, whereas *At*GORK^623^ has strict C4 symmetry. The displacement of Cα position of Tyr196 in the C4-C5 region is approximately 3.8 Å (indicated by yellow dash-line). (D) further compares protomers A and B of both *At*GORK^FL1^ and *At*GORK^623^, showing that the protomers in *At*GORK^623^ closely resembles protomer A in *At*GORK^FL1^, rather than protomer B. ANK truncation in *At*GORK^623^ restores C4 symmetry in the tetrameric assemblies.

We also performed AlphaFold3-based modeling on *At*GORK^623^, showing that the TMD of the predicted *At*GORK^623^ structure is highly similar to that obtained by cryo-EM, with an RMSD of 1.97 Å for the superimposed tetramers. However, the subsequent portions, including the C-linker, CNBD, and ANK domains, are in a loose and relaxed conformation. Due to the lack of ANK dimerization, the overall structure exhibits C4 symmetry (Figure S8, Supporting Information). Together with our structural and electrophysiological data, these observations support the role of ANK as a molecular switch for symmetry conversion in GORK, revealing a novel mechanism for channel regulation via symmetry conversions.

### A model for symmetry conversion in GORK regulation

In summary, our analysis shows that full-length *At*GORK^FL^ adopts C2 symmetry due to ANK dimerization in the cytoplasm. Cryo-EM and AlphaFold3 modeling of ANK-truncated *At*GORK^623^ reveal a restoration of C4 symmetry within the tetrameric assembly. This symmetry conversion, driven by ANK dimerization, regulates intermodular communication and channel gating process. The transition between C2 and C4 symmetry governs GORK switching between autoinhibited and activatable states. In short, ANK dimerization maintains GORK in C2 symmetry and an autoinhibited state, while disrupting ANK dimerization restores C4 symmetry and activates GORK (Figure 6). Thus, the ANK domain acts as a molecular switch for symmetry conversion, providing a finely tuned mechanism for GORK regulation.

**Figure 6.**
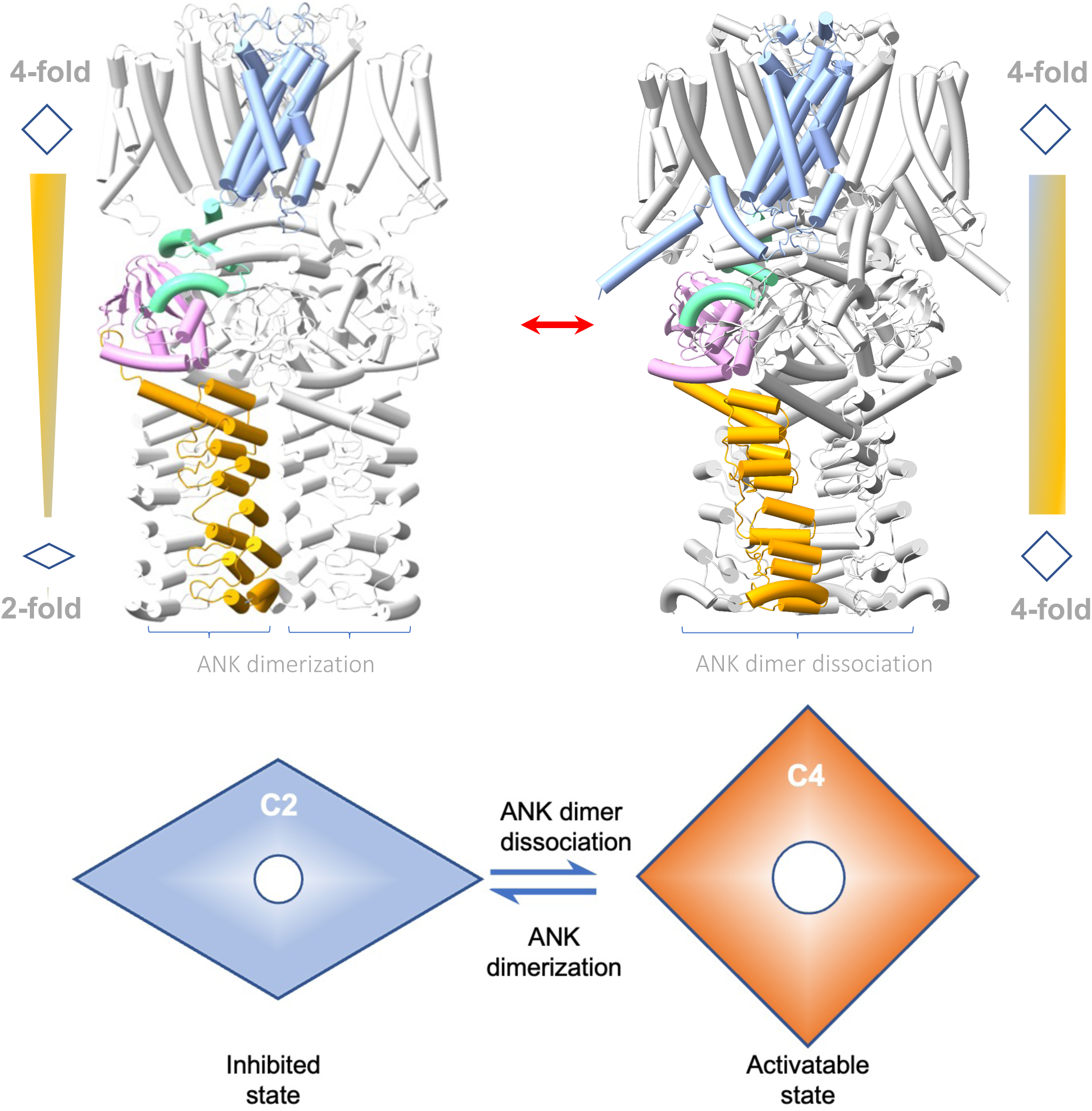
A model of symmetry conversion in plant GORK regulation. The GORK structure in the autoinhibited state (left, Cryo-EM) exhibits C2 symmetry, while in the activatable state (right, Alphafold3 prediction), it exhibits C4 symmetry. In brief, ANK dimerization reduces the symmetry of the GORK tetramer from C4 to C2, resulting in an autoinhibited state. Disrupting dimerization restores C4 symmetry, converting GORK to an activatable state. This dynamic symmetry conversion in the GORK tetramer provides a unique mechanism for regulating channel activity, enabling transitions between inhibited and activatable states - a distinctive feature of GORK family proteins containing the ANK domain. In such a way, ANK functions as a molecular switch, regulating the GORK channel in guard cells and controlling stomatal movement in response to environmental stimuli.

## Discussion

Plant potassium channels are crucial for K^+^ transport and maintaining potassium homeostasis^[2]^. In *Arabidopsis*, nine Shaker-like potassium channels contain multiple modular domains in their architectures, including the pore-forming TMD, the regulatory C-linker, and CNBD, with some also containing ANK and KHA domains. Based on the presence of the ANK domain, these channels can be divided into two groups: ANK-free members (KAT1, KAT2, and KC1) and ANK-containing members (GORK, SKOR, AKT1, AKT2, AKT5, and AKT6). Currently, structural information are available for KAT1^[20–21]^, SKOR^[26]^, and AKT1^[27,31]^, allowing us to conduct systematic comparative analysis

In ANK-free KAT1, structural studies show that it forms a symmetric tetramer with typical C4 symmetry throughout both the transmembrane and intracellular regions^[20–21]^, similar to animal cyclic nucleotide-gated channels (CNGC)^[30]^. In contrast, GORK, which contains a unique ANK domain, exhibits a symmetry transition from C4 in the transmembrane region to C2 in the cytoplasmic region, resulting in distinct structural features. A similar symmetry reduction is observed in other ANK-containing potassium channels, such as AKT1^[27]^ and SKOR^[26]^. This reduction in symmetry is rare among ion channels, suggesting it may be a unique feature of ANK-containing channels linked to their regulatory mechanisms.

Our study provides the first clear-cut evidence that ANK acts as a molecular switch controlling GORK channel symmetry and activity. Structural analysis reveals that ANK dimerization holds GORK in an autoinhibited state at rest. ANK dimerization mediates interactions between protomers, reduces tetramer symmetry, and causes the displacement of internal, highly mobile modules, with neighboring CNBDH and C-linker shifting from C4 symmetry to C2. The C-linker adopts two different conformations: “kinked” and “flat” (Figure 2B). The “kinked” conformation is common in C4 symmetric channels, while the “flat” conformation is unique to ANK-containing channels with reduced symmetry. Subtle yet distinct differences in the S5 helices between adjacent protomers in *At*GORK^FL1^ renders a quasi-C4 symmetry in the TMD (Figure 5B). The reduction in GORK symmetry signifies an autoinhibited state, supported by the observed basal electrophysiological activity.

Our ANK truncation experiments showed that ANK removal converts GORK from an autoinhibited to an activatable state, as evidenced by enhanced activities in the truncated *At*GORK^543^ and *At*GORK^623^ variants upon membrane depolarization (Figure 3B-E). Structural analysis further revealed that the TMD restores strict C4 symmetry, while the C-linker and CNBDH become highly mobile in these ANK-truncated mutants. Additionally, AlphaFold3-based modeling confirmed that disrupting ANK dimerization relieves constraints and restores C4 symmetry in the tetramer, enabling GORK transition to an activatable state. Thus, ANK serves as a molecular switch to regulate GORK activity by controlling the symmetry conversion of its tetrameric assembly.

Ankyrin domains also occur in TRPV channels, where they form a “petal-like” arrangement in the cytoplasm and act independently rather than directly participating in channel assembly^[32]^. Typically, these domains consist of six ankyrin repeats, forming a “palm-and-finger” structure, whose concave surface binds cytoplasmic proteins or ligands. Unlike GORK, and TRPV gating depends on ANK conformational changes, such as rigid-body rotation. In TRPV4, RhoA binding to the ANK domain inhibits channel activity, whereas calmodulin binding enhances it, highlighting the versatile roles of ANKs in fine-tuning channel activity and coordinating intracellular signaling.

The autoinhibited-to-activatable transition of the GORK channel is governed by the dimerization state of its ANK domain, yet the molecular triggers of this dimerization and its reversal remain unknown. A recent study suggests that the cytosolic N-terminus of GORK functions as a “safety catch”, constraining its transitions toward the open state^[33]^. In the truncated mutant GORK^Δ23^, removal of the N-terminal segment not only relieves autoinhibition and markedly enhances channel activity, but also restores C4 symmetry in the pre-opened conformation. This provides additional evidence for a C2– C4 symmetry switch during channel assembly. Thus, both the N- and C-termini are crucial for GORK regulation, but how they cooperate is still to be elucidated.

Previous studies showed that phosphorylation affects ANK-mediated protein-protein interactions and channel activity^[32,34–35]^. Several protein kinases, including CPK21^[36]^ and CPK33^[37]^, have been identified as candidates for regulating GORK activity, with phosphorylation sites, such as S649 in the ANK domain^[38]^. However, the molecular mechanisms of these kinases require further investigation. A parallel case is AKT1, which is activated by CIPK23 and CBL1, with multiple potential phosphorylation sites identified (including within its ANK domain). The low-resolution structure of AKT1 co-expressed with CIPK23 and CBL1 shows C4 symmetry in the CNBDs, though it remains unclear whether this is due to ANK phosphorylation^[27]^.

Autoinhibition is an intrinsic mechanism that maintains ion channels and transporters in an inactive state, conserving energy in the resting state. In plants, different proteins achieve this through distinct molecular strategies. For example, SLAC1—a guard-cell anion channel essential for stomatal closure—is held in its autoinhibited state by interactions between its N-terminal domain and the pore-forming transmembrane region; phosphorylation relieves this inhibition, rendering the channel activatable^[39]^. By contrast, GORK utilizes its ANK domain as a molecular switch: ANK dimerization reduces the tetramer symmetry (from C4 to C2) to ensure autoinhibition, while disrupting ANK dimerization restores C4 symmetry and allows GORK becoming activatable. Together, these cases highlight the diversity of auto-inhibition mechanisms and shed light on their roles in plant physiology.

In summary, our study reveals that ANK regulates GORK channel activity through tetrameric symmetry conversion. Under resting conditions, ANK dimerization maintains GORK in a C2 symmetric conformation, maintaining it in a preactivated, autoinhibited state. In response to environmental stimuli, GORK undergoes a symmetry conversion and runs into an activatable state, enabling K^+^ efflux and stomatal closure. This mechanism provides new molecular insights into stomatal regulation and offers potential avenues for developing improved crop varieties through ANK engineering.

## Methods

### Expression and purification of the GORK proteins

The genes of *Arabidopsis* full-length *At*GORK (AT5G37500) and truncated variants (*At*GORK^623^, residues 1-623 and *At*GORK^510^, residues 1-510) were cloned into a modified pEG BacMam vector, which included a PreScission protease cleavage site, GFP, an 8× His tag, twin-Strep tag, and Flag tag at the N-terminus. Recombinant proteins were expressed in HEK293F cells. In short, baculovirus was generated in *Sf9* insect cells, and P2 viruses were used to infect HEK293F cells. Cells were cultured in suspension, supplemented with 1% (v/v) fetal bovine serum, and maintained at 37 °C with 70% humidity and 6% CO_2_. Cells were infected with P2 viruses at a density of approximately 2.6×10^6^ cells/mL, and supplemented with 10 mM sodium butyrate upon 8 hours post-infection, and continued to grow at 30℃ for another 50 hours. Cells were harvested, centrifuged, and stored at −80°C.

Harvested cells from a 300 mL of culture were resuspended in extraction buffer (20 mM HEPES-Na pH 8.0, 150 mM KCl, 10 mM DDM, 2 mM cholesteryl hemisuccinate (CHS) supplemented with protease inhibitors (2 μg/mL pepstatin, 2 μg/mL aprotinin, 2 μg/mL leupeptin and 1mM phenylmethylsulfonyl fluoride) and incubated at 4℃ for 2 hours. Solubilized membranes were clarified by centrifugation at 41,000 rpm for 1 hour, and the supernatant was loaded onto a Streptactin Beads 4FF column. Following a 5-column volume buffer wash, proteins were eluted with solubilization buffer (20 mM HEPES-Na pH 8.0, 150 mM KCl, 0.02% GDN, and 2.5 mM desthiobiotin). After tag removal with TEV protease, the proteins were further purified using a Superose-6 column in solubilization buffer (20 mM HEPES-Na pH 8.0, 150 mM KCl, 0.006% GDN). The elution was analyzed by SDS-PAGE, and the peak fractions of the purified protein were pooled and concentrated to ∼3.5 mg/mL.

For the preparation of truncated truncated versions (*At*GORK^623^ and *At*GORK^510^), we followed similar procedures as above.

### Cryo-EM grid preparation and data acquisition

The protein sample (4 μl, ∼3.5 mg/ml) was applied to freshly glow-discharged holy carbon film grids (Quantifoil, Cu, R1.2/1.3, 300 mesh) and blotted for 6.5 seconds at 100% humidity and 4°C using a Vitrobot Mark IV (Thermo Fisher Scientific), followed by plunge freezing into liquid ethane.

All the films were collected on a Titan Krios transmission electron microscope (Thermo Fisher Scientific, USA) operated at 300 kV equipped with a Gatan K2 or K3 Summit direct detection camera (Gatan Company, USA). Movies were recorded at 22,500× (for K3, corresponding 1.06 Å per pixel) or 130,000 × (for K2 summit, corresponding 1.04 Å per pixel) nominal magnifications, using SerialEM software with a beam-image shift method^[40–41]^, at a total dose of 50 e-/Å2 distributed over 32 frames, with defocus values between −1.2 and −2.2 μm.

### Data processing and model building

All image processing steps were conducted using either CryoSPARC 3.1^[42]^ or RELION 3.0^[43]^. CryoSPARC 3.1 was primarily used for reconstruction, while RELION 3.0 was employed for alignment-free 3D classification to identify an alternate conformation (*At*GORK^FL2^) of the wild-type structure. Angular information was converted using PyEM, and map analysis and adjustments were performed with UCSF Chimera^[44]^.

Raw movie frames were aligned using a dose-weighted patch alignment method, followed by CTF estimation. Particle picking was performed an oval template (88 × 168 Å), and coordinates were screened using normalized cross-correlation and local power scores to eliminate false positives. Particles were then down-sampled to a quarter size and classified based on shape (circular for top-view projections and T-shaped for side-view projections) after initial 2D classification to improve classification efficacy. A preliminary 3D reconstruction was generated from a small subset of particles.

*At*GORK^FL1^ was constructed with Nu-refinement using C2 symmetry with 156,313 particles, while *At*GORK^FL2^ used 39,551 particles with C1 symmetry. The resolutions were 3.4 Å for *At*GORK^FL1^ and 4.3 Å for *At*GORK^FL2^, respectively. The alphafold2-predicted *Arabidopsis* GORK structure was used as the initial model, fitted by individual domains in UCSF ChimeraX^[45]^, and refined iteratively in Coot^[46]^ and PHENIX^[47]^, and corrected iteratively. For the truncated *At*GORK^623^ and *At*GORK^510^, the TMD structure from *At*GORK^FL1^ was used as the initial model, followed by iterative refinement and iterative adjustments in Coot and PHENIX. The resolutions were 3.2 Å for *At*GORK^623^ and 3.4 Å for *At*GORK^510^, respectively. All figures were generated using UCSF ChimeraX.

### Electrophysiology

The electrophysiological experiments were performed as previously described^[48]^. Briefly, untagged coding sequences of *At*GORK wild type or mutated variants were cloned into the pGHME2 vector for expression in *Xenopus* oocytes. cRNAs were synthesized using T7 polymerase with linearized plasmid DNA templates. Oocyte sacs were extracted and digested with 0.2 mg/ml collagenase (Sigma) in OR2 buffer (82.5 mM NaCl, 2.5 mM KCl, 1 mM MgCl_2_, and 5 mM HEPES-Na pH 7.5). For expression, 36 ng of wild-type GORK cRNA was injected per oocyte, with mutant cRNA doses adjusted to equimolar concentrations. Injected oocytes were incubated at 18°C in ND96 buffer (96 mM NaCl, 1.8 mM CaCl_2_, 1 mM MgCl_2_, 2 mM KCl, and 5 mM HEPES-Na pH 7.5) for 48 hours before use.

TEVC recordings were performed using an OC-725C amplifier (Warner Instruments) and Digidata 1550B (Molecular Devices) controlled by pClamp software. The bath solution contained (in mM): 10 KCl, 90 NaCl, 1 CaCl₂, 2 MgCl₂, and 10 Tris/MES (pH 7.4), with osmolarity adjusted to ∼220 mOsmol/kg using D-mannitol. Microelectrodes (0.5-1 MΩ resistance) were filled with 3 M KCl, and agar bridges served as bath electrodes. Voltage steps from −110 mV to +70 mV (20-mV increments, 2,000 ms duration) were applied, followed by −110 mV clamping for 450 ms, 4,000 ms, or 16,000 ms depending on deactivation kinetics. Steady-state currents were measured at 50 ms before the end of each step. Data were analyzed using Clampfit 10.6 (Molecular Devices), Prism 8.0 (GraphPad), and Origin 2021.

### Fluorescence and confocal imaging

To assess the cell-surface expression of full-length and C-terminal truncated GORK channels in *Xenopus* oocytes, GFP was fused to the N-terminus of each construct. The corresponding cRNAs were injected into oocytes, which were then incubated at 18°C for ∼48 hours in ND96 buffer (detailed in the *Electrophysiology* section). GFP-tagged protein expression was verified using a confocal laser scanning microscope (Zeiss LSM 980), and fluorescence intensities were quantified with ImageJ. To minimize variability in expression levels due to experimental factors (e.g., cRNA injection dose or incubation time), all experiments were performed in parallel.

## Supporting information

suppFigures

## Data availability

All data generated or analysed in this paper are presented in the main text, figures and the extended data figures and supplementary videos, or are available from the corresponding author upon request. The cryo-EM maps of the *At*GORK full-length (*At*GORK^FL1^ and *At*GORK^FL2^) and truncated version (*A*GORK^623^ and *A*GORK^510^) have been deposited in the Electron Microscopy Data Bank with accession codes EMD-62338, EMD-37500, EMD-62337 and EMD-62339, respectively, and their structural models have been deposited in the PDB with accession codes 9KHF, 8WFZ, 9KHE and 9KHG, respectively (Table S1, Supporting Information).

## Acknowledgments

This project is financially supported by the National Key Research and Development Program of China (2020YFA0509903, 2016YFA0500503 and 2021YFA1300702), the National Natural Science Foundation of China (32300257, 31322005 and 31470728), and The Chinese Academy of Sciences Project for Young Scientists in Basic Research (YSRB-119).

## Author contributions

Q.-Y.L. performed protein purification, Cryo-EM data collection, data analysis and mutant design; L.-H.T. and C.-R.Z performed Cryo-EM data collection, structural determination and analysis; L.Q. performed electrophysiology and data analysis, K.-W. and G.-H.Z. performed experiments; S.-G.H., Q.X., T.-X.N. and M.S. analyzed data; R.H., analyzed data and wrote the manuscript, Y.-H.C. initiated the project, planned and analyzed experiments, supervised the research, and wrote the manuscript with input from all authors.

