## Supplementary material for "Structural and Mechanistic Insights into Symmetry Conversion in Plant GORK K^+^ Channel Regulation": suppFigures

### Figure S1

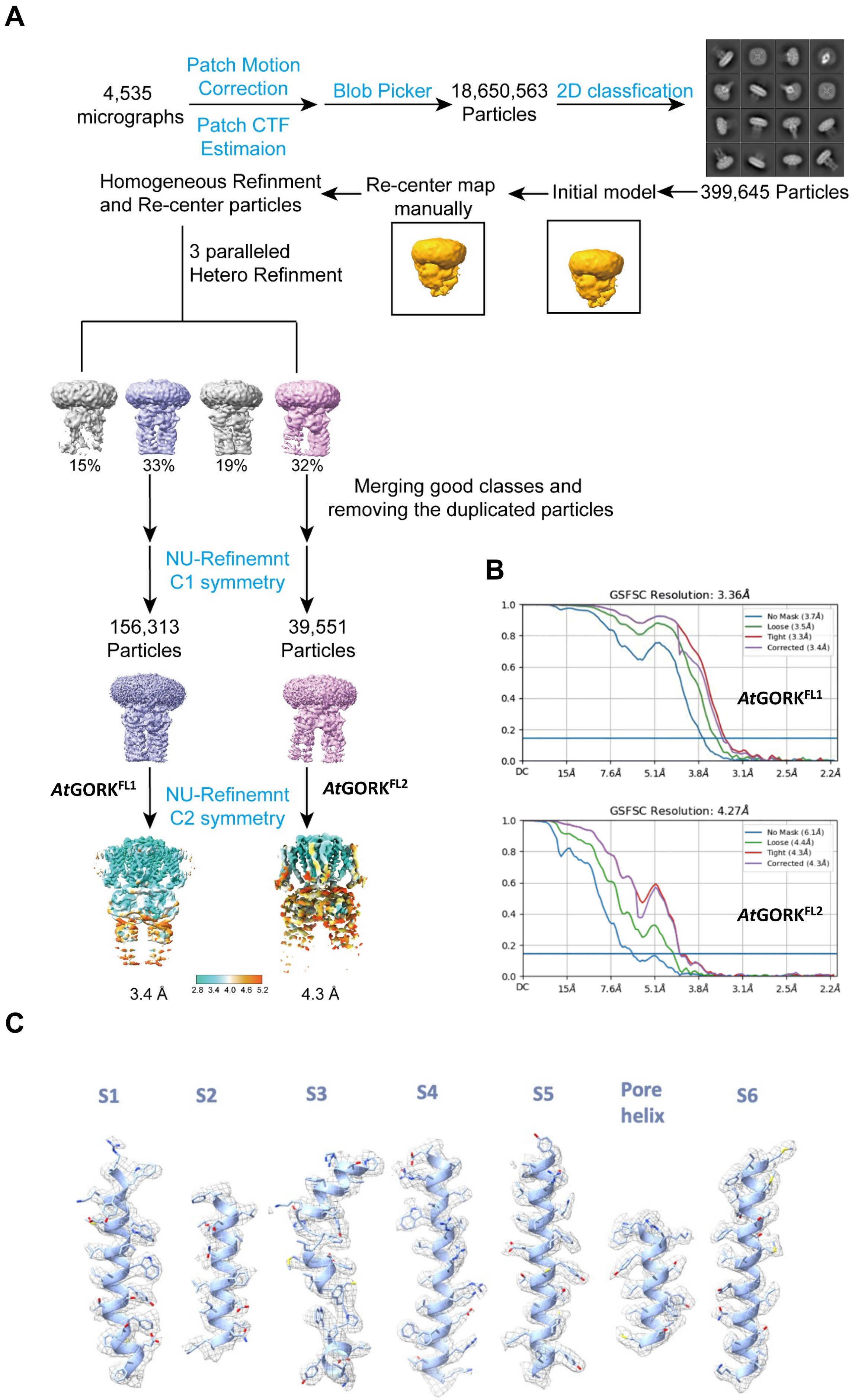
A
4,535
micrographs
Patch Motion
Correction
Patch CTF
Estimaion
Blob Picker
18,650,563
Particles
2D classfication
Homogeneous Refinment
and Re-center particles
Re-center map
manually
Initial model
399,645 Particles
3 paralleled
Hetero Refinment
15%
33%
19%
32%
Merging good classes and
removing the duplicated particles
NU-Refinemnt
C1 symmetry
156,313
Particles
39,551
Particles
AtGORKFL1
NU-Refinemnt
C2 symmetry
AtGORKFL2
3.4 Å
2.8 3.4 4.0 4.6 5.2
4.3 Å
B
GSFSC Resolution: 3.36Å
No Mask (3.7Å)
Loose (3.5Å)
Tight (3.3Å)
Corrected (3.4Å)
AtGORKFL1
GSFSC Resolution: 4.27Å
No Mask (6.1Å)
Loose (4.4Å)
Tight (4.3Å)
Corrected (4.3Å)
AtGORKFL2
DC
15Å
7.6Å
5.1Å
3.8Å
3.1Å
2.5Å
2.2Å
C
S1
S2
S3
S4
S5
Pore
helix
S6

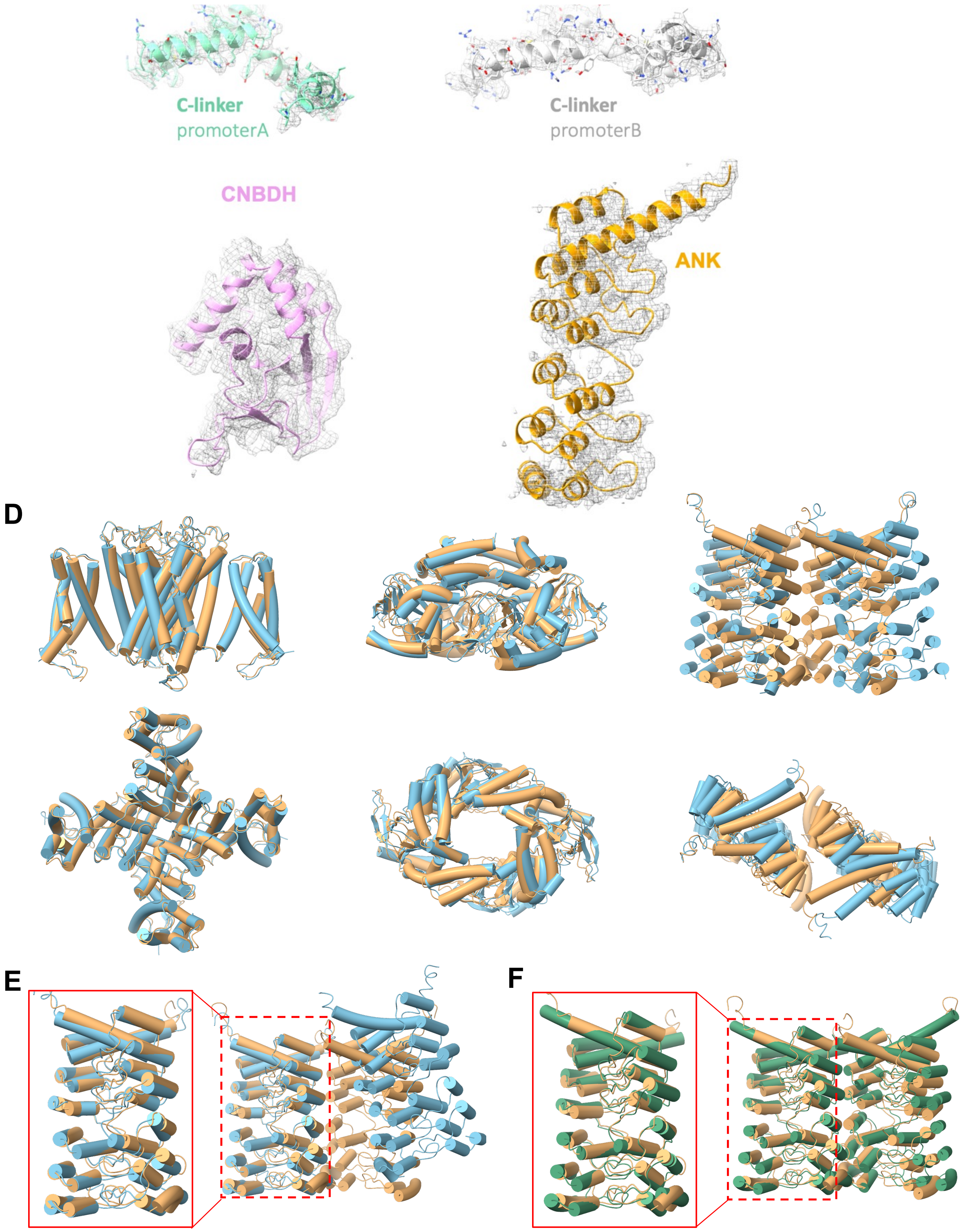


**Figure S1. Structural determination of full-length *Arabidopsis* GORK (*At*GORK$^{FL}$).**

(**A**) Workflow for image processing of full-length *Arabidopsis* GORK. (**B**) Fourier shell correlation (FSC) curve indicating overall resolution of 3.4 Å for *At*GORK$^{FL1}$ and 4.3 Å for *At*GORK$^{FL2}$, as estimated using the 0.143 cut-off criterion (dotted line). (**C**) Representative cryo-EM density map showing TM segments (S1-S6) and domains (C-linker, CNBDH and ANK). (**D**) Structural comparison of domains in *At*GORK$^{FL1}$ (light orange) and *At*GORK$^{FL2}$ (light blue). (**E**) Superimposition of ANK dimers in *At*GORK$^{FL1}$ (light orange) and *At*GORK$^{FL2}$ (light blue), with a RMSD of 1.21 Å/dimer. (**F**) Superimposition of ANK dimers in *At*GORK$^{FL1}$ (light orange) and *At*SKORK (purple, 8jet), with a RMSD of 1.33 Å/dimer. Despite variations in ANK conformation across the tetramer, the dimeric interaction is conserved, indicating that ANK dimerization is a fundamental structural feature of potassium channels with ANK domains.

### Figure S2

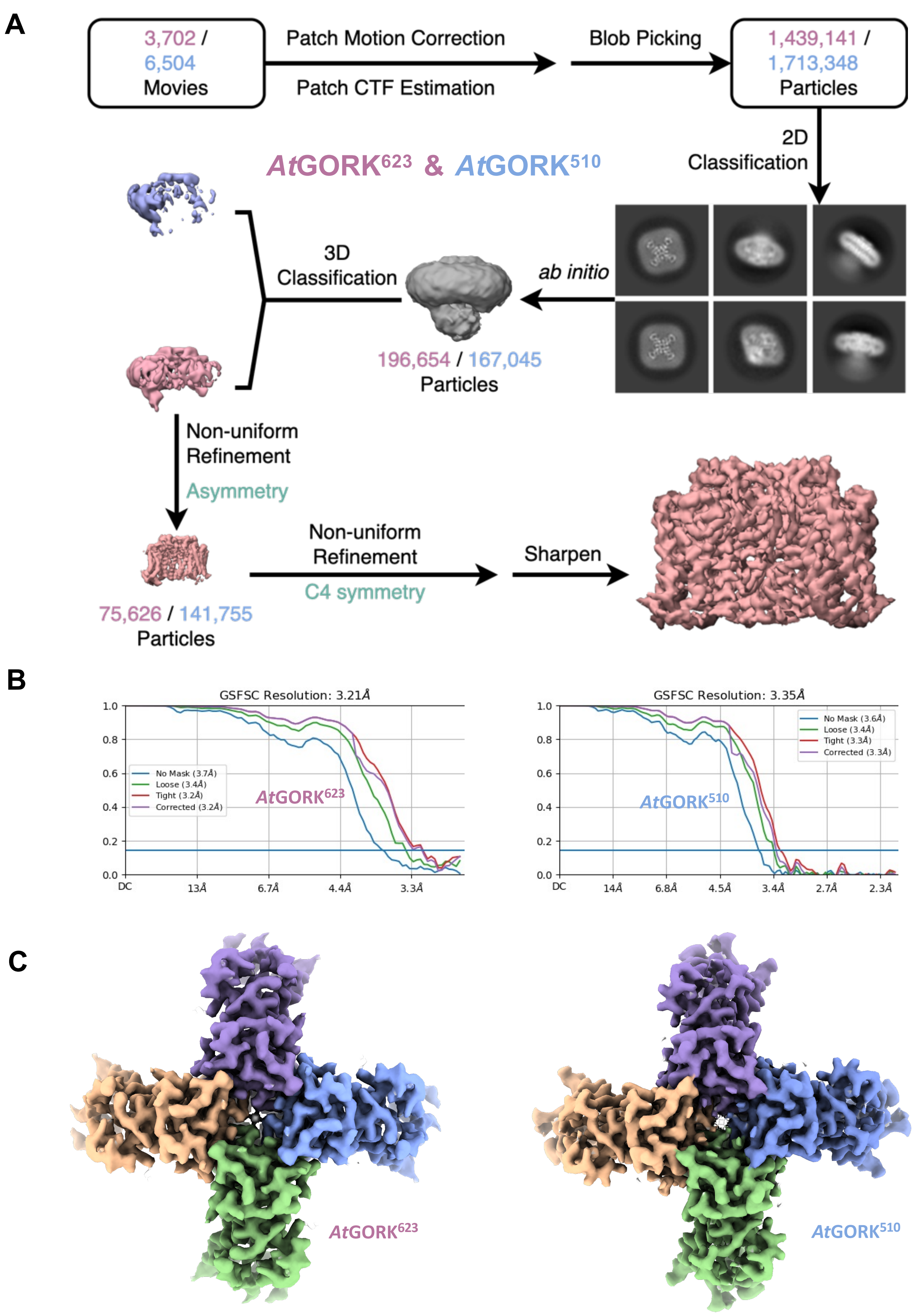


**Figure S2. Structural determination of the truncated *At*GORK$^{623}$ and *At*GORK$^{510}$.**

(**A**) Workflow for image processing of the truncated *At*GORK$^{623}$ (residues 1-623) and *At*GORK$^{510}$ (residues 1-510). (**B**) Fourier shell correlation (FSC) curve indicating overall resolution of 3.2 Å for *At*GORK$^{623}$ and 3.4 Å for *At*GORK$^{510}$, as estimated using the 0.143 cut-off criterion (dotted line). (**C**) Cryo-EM structures of *At*GORK$^{623}$ (3.2 Å, left) and *At*GORK$^{510}$ (3.4 Å, right).

```
                                                                                          S1
Arabidopsis_GORK     --------------MGRL-----RR---R-------QEI-------IDHEEEESNDDVSSRRGKLSLAETFRWLDSSEHRRIET--------DGH--NDYKYIIHPK-NRWYKAWEMFIL   73
Brassica GORK        --------------MGCL-----RR---R-------QES-------IAE-EDDINDDVSRRRGRFSLAESFRWLDSPEHLKDDS--------DGH--NEYPWIIKPS-IRWYKAWELFIL   72
Zingiber_GORK        MERDSSRRK-WKRTMSAVMAPRPSEAEEEAVGLASDGGSGGGDEYTVEEVHDLIG---SSRGSRQSVLADESRR-----------------ASGDELLLRFVFRPD-NGWYETWTQFIM   97
Elaeis GORK          -------------------------------MSPRAKNGSPEIEYEL---EEPIA---SSRGIRLFLLTSEFALGPLRRRAT----------SQEKLLERFVIEPD-NRWYQLWTRFIL   72
Phoenix_GORK         -------------------------------MSPHAKNESQEIEYEL---EEPMG---SSRGIRWSLITSDFAIGPLRRR-AH----------SHEKLLERFVIEPD-NKWYQLWTRFIL   71
Oryza GORK           MGRGIGSKRRVEDDDGENMPGRK---KKE--EEEEEEDDDGEEEYEVDVVRDRIG---SSRGSRLALFGSDLRLGRFRPRRRRV--APV---DGDDGIFQDFVIDPD-NKWYRLWTRFIL  106
Brachypodium_GORK    MGRGLGSKSRVES-------------------------EVEEEEYEVEVVPDRLK---SSRNSRLALFGSELRLDRFRPRRRRRRAAA---DGEDGFFHDLIIHPE-NKWYRIWSRFIL   88
Panicum GORK         MGRGLGSKRRVAEGDTK----------------PEEEEEGSSEEYEVEVVRDHIA---SSRGSRLALFGSDLRLGRFRPRRRRA---------AAEGFFQDLVIHPD-NRWYRLWTKFIL   91
Papaver_GORK         MPD----------HLHHL-----HRHHHERERDKREEDEDETEEFKVEKLKDRIL---SSRGSRFDLITNELGLDNQ-TRRKISREDFI---NGIR----GFVIHPD-NRWYLIWTRIIL   93
Amborella GORK       -------------------------------MCPGSEERDMQEEYQVEEFRERIN---SSRGSRFALLSELTKR---------GGGSQERLIQGFRGCTQSVIIHPD-SRWYQVWTFIIL   76
Vulgaris_GORK        ---------MTPKQLGRIISGKTMSRIREEGGSTSSSSSNRIEDFVVEKPRERLT---SSRGSRLDLLVNELNDES--TRRQFSREGFF---LGVKDLYHGLVIHPEDNRWYKAWDKFIL  103
Chenopodium GORK     ---------MKPKQIARIMMTRG--REERMEGSTSSSSSNREEDFMVERPRERLY---ASVGSRFNLLVTEFSHES--SQRQFTREGFF---VGVKDLYHGLVIHPEHNRWYKAWDNIIL  101
Lactuca_GORK         ------------MRIGST---------ADERRRKAVEDEVVGEEYVVEDVRDHDHGRLTKRRSRFTLLAGDLDLDP--SRRDFNTEK---FIDGIRGCSPALFIHPQ-NSWYKAWEKFIL   93
Daucus GORK          ----------------MSNRRRMEEEKRRRRKGQDSEDSSNDEEYEVEDVREDVR---ESRGHHMARIASEFEAATEAYRRIYSREHIF---KGLTDLSRGCSIHPD-NRWYHAWEKFIL   97
Medicago_GORK        ---M---KMKEEEDEGEKNRINNQTATCSGSSSLTPSSSSDEREYQVQDLRDKLK---SSRGSRFNLIEKELGLKI--GWRKFSR----------RTLLHEFVINPL-NRWYRAWLKFIL   98
Prosopis GORK        MPGN---RLTEEAVTEEREEKRKQMLSTSRSSSSSSSSSSDDREYELEDLRDRLK---SSRGSRFQLIENELKE-L--NQKKFSRHV----IDGIRGLSKDFIIHPE-NRWYRAWTKFIL  106
Nicotiana_GORK       -------------------------MTRVAEEVLETMNMNGGVEYEVEDVRDKIE---SSRGSRFQLIEDDL-GLNL-FRRRISRL--------KINLSQVFIIHPE-NRWYRMWENFIL   81
Solanum GORK         --------------------------MTRGEEVLEIDINGGRAEYEIEDMRDKME---SSRGSRFKLIENDLVGADM-FSRRRRKFSRESLLNGLKDLSQGFVIHPE-NRWYRMWENFIL   89
Coffea_GORK          -------------------------MGRGNSHREEGESRNGEGEYEVEDLRDRIK---SSRGSRFELIEHELGL-DS-THRRFSRQSVI---NGLKDLSQGLVIHPD-NRWYRAWEKFIL   86
Sesamum GORK         --MG---SFKKSSF--RN-----TDESSANGEESSLNGVNGGEEYMVEDLRERIK---SSRGSRFTLIENELGLIDS-ARRRVSRQTVI---DGLKDLSQGLFIHPD-NRWYRAWEKFIL  100
Cucumis_GORK         -----------MAVSG-------GREKEEEAGTHRRSDSDGEVEYRVESVRERIS---SSRGSRFNLIQRQLRID--QSRRRFSRENLI---NGIK----CLVILPD-SRWYRAWTKFIL   89
Vitis GORK           --MN---RFMY--LVGRG----------------QVVDDEDDGDFQMPDLRERIK---GSSASVLNMFNGDLGLR------RFSGGSCF---NGIK----GFIILPD-NWWYRTWTKFIL   80
Cannabis_GORK        ---M---SFAAANGIGRV-----DQNDERDHDMVSSSSDGDEEEFELQDIRDRIK---SSRGSRFNLIANEFGLASAI--KKFSRENVM---NGIRDLSNGVVIHPD-NRWYRAWTKFIL  100
Gossypium GORK       -----------MGDIGRS-----RRV-----RDERCATMANEEEYEVEDFREGNK---SSRGSRFNLIANELGLVAARARMNLSRQSVL---NGIKDLSDGP-IHPD-NRWYRAWTKFIL   91
                                                                 :    :                                                  : *     **  *  :*:
```

```
                         S1                          S2                           S3a        S3b      S3c                  S4
Arabidopsis_GORK     VWAIYSSLFTPMEFGFFRGLPE--RLFVLDIVGQIAFLVDIVLQFFVAYRDTQTYRTVYKPTRIAFRYLKSHFLMDFIGCFPWDLIYKASGKHELVRYLLWIRLFRVRKVVEFFQRLEKD  191
Brassica GORK        VWAIYSSLFTPMEFGFFRGLPE--NLFILDIVGQIAFLVDIVLQFFVAFQDKHTYRIDSKPTHIALRYLKSHFFLDLVSCFPWDLIYKASGKHEVVRYILWIRLFRVRKVIEFFQRLEKD  190
Zingiber_GORK        LWAVYSSFFTPLEFGFFRGLPK--NLIFLDMTGQVAFLVDIIVNFFLAYRDSLTYRMIYSPTSIAFRYLKSSFVFDFLACVPWDYIYKLSGRKEEVRYLLWIRLIRVRKVTDFFQRMEKD  215
Elaeis_GORK          VWALYSSFFTPIEFGFFRGLPN--NLFWLDFAGQVAFLVDIFVQFLVAYRDSHTYRMIYEPTSIAVRYAKSSFVFDLLGCFPWDAIYRACGRKEEVRYLLWIRLTRVRKVTDFFQKMEKD  190
Phoenix_GORK         VWALYSSFFTPIEFGFFRGLPN--NLFWLDFAGQVAFLVDIFVQFLVAYRDSHTYRMIYKPTSIAVRYVKSSFVFDLLGCFPWDVIYKACGNKEEVRYLLWIRLTRVRKVTDFFQKMEKD  189
Oryza_GORK           VWAVYSSFFTPLEFGFFRGLPR--NLFFLDIAGQIAFLIDIVLRFFVAYRDPDTYRMVHNPTSIALRYCKSSFIFDLLGCFPWDAIYKACGSKEEVRYLLWIRLTRAMKVTEFFRSMEKD  224
Brachypodium_GORK    VWAVYSSFFTPFEFGFFRGLPK--RLFFLDIAGQIAFLIDIVLKFFVAYRDPDTYRMVYNPTSIALRYCKSSFIFDLLGCFPWDVIYKACGSREEVRSLLWIRLTRALKVTEFFKDLEKD  206
Panicum_GORK         VWAVYSSFFTPLEFAFFRGLPR--KLFLLDIAGQIAFLIDIVVKFFVAYRDPDTYRIVYDPTAIALRYFKSSFIFDLLGCFPWDVIYKACGRKEEVRYLLWIRLTRALKVTEFFWHLEKD  209
Papaver_GORK         IWAIYSSFFTPVEFGFFRGLPD--KLFILDIAGQFAFFVDIIIQFFVAFRDPQTYRMVSKRTTIALRYLKSSFILDLLACCPWDLIYASCGKKEEVRYLLWIRLYRVRKVIEFFQHMEKD  211
Amborella_GORK       MWAIYSSFFTPLEFGFFRGLPK--NLVFLEVAGQIAFLVDILFNFFLAYRDSQTYRMVYKRSDIALRYAKSCFLVDLFGCLPWDAIYKASGRKEVVRYLLWVRLSRLQKVTNFFQRLEKD  194
Vulgaris_GORK        LWAIYSSFFTPMEFGFFRGLPE--NLFVLDIVGQVAFLLDIVLQFFVGYRDKQTYRMVYQRPAIAFRYLKSTFVIDLLACMPWDLIYKASHHKEAVRYLLWIRLCRVRKVHYFLQKMEKD  221
Chenopodium_GORK     LWAIYSSFFTPMEFGFFRGLPE--NLFVLDIVGQVAFLMDIVLQFFVGYRDKQTYRMVYQRPAIAFRYLKSTFVLDLLACLPWDLIYKASHRKEAVRYMLWIRLCRVRKIHFFLQKLEKD  219
Lactuca_GORK         IWAVYSSFFTPMEFGFFRGLPK--NLYFLDIAGQIAFLIDVVLHFFIAYRDTQTYKMISNRNLIAVRYLKSHFLFDILACMPWDNIYRASGRKEEVRYLLLLRLVRVRKVLEFFSRLEKD  211
Daucus_GORK          LWAMYSSFFTPFEFGFFRGLPRNRNLFLLDIAGQTAFVIDIVLQFFVAYRDRQTYKMIYKRYPIAMRYLKSHFIIDFLGCLPWDIIYGATGNKEAVRCLLWIRLSRARKVLAFFQKLEKD  217
Medicago_GORK        LWAVYSSFFTPMEFGFFRGLPE--NLFILDIVGQIAFLVDIVLQFFVAYRDSQTYRMVYKRTPIALRYLKSTFVIDLLGCMPWDLIYKACGRREEVRYLLWIRLYRAERVVQFFRNLEKD  216
Prosopis_GORK        LWAIYSSFFTPMEFGFFRGLDE--DLFILDIIGQVAFLVDIIVQFFLAYRDSQTYRMIYKRAPIALRYLKSSFIFDLLGCMPWDIIYKASGRREGVRYLLWIRLYRVRKVTEFFQRLEKD  224
Nicotiana_GORK       IWSIYSSFFTPMEFAFFNGLPR--KLFLLDICGQIVFLVDIVLQFFVAYRDSQTYKMVYKRTPIALRYLKSHFILDFLSCMPWDIIYKAVGSKEEVRYLLWIRLSRARRITYFFQKMEKD  199
Solanum_GORK         IWSIYSSFFTPMEFAFFNGLPR--KLFLLDICGQIVFLVDIVIQFSVAYRDSQTYKMVYKRTPIALRYLKSHFIMDFLGCMPWDIIYKAVGSKEEVRYLLWIRLSRARRITYFFQKMEKD  207
Coffea_GORK          IWAVYSSFFTPFEFGFFRGLPR--KLLFMDIAGQAAFLVDIILQFLVAYRDSQTYRMVYKRTPIALRYIKSHFIIDLLGCMPWDFIYKAVGHKEVVRYFLWIRLSRVRKGNKFFQHMEKD  204
Sesamum_GORK         IWAIYSSFFTPMEFGFFRGLPK--NLFFLDIAGQVAFLLDIILQFFVTYRDSHSYKMVYNRNLIALRYLKSHFLPDLLGCMPWDIIYKAVGEKEEVRYLLWIRLTRVRKVTAFFQRMEKD  218
Cucumis_GORK         IWAVYSSFFTPMEFGFFRGLPE--NLFILDIVGQIAFLFDIVFQFFLAYRDKQTYRMVYKRSPIALKYLKSTFVTDLLSCMPWDILYKACGRREEVRYLLWIRLFRVRKVDAFFKTMEKD  207
Vitis_GORK           LWAVYSSFFTPMEFGFFRGLPE--DLVFLDIAGQIAFLIDIVLRFFLAYRDAHTYRMVYKRTSIALRYMKSSFVIDLICCLPWDIIYKACGRKEEVRYLLWIRLIRVCKVTDFFQNLEKD  198
Cannabis_GORK        LWAVYSSFFTPMEFGFFRGLNE--DLFVLDIVGQIAFLVDIVVLFFVSYRDSHTYRMVYKRTPIALRYLKSSFVIDLLCCLPWDIIYKKSGRHEAVRYLLWIRLSRVRKVTDFFHKLEKD  218
Gossypium_GORK       IWALYSSFFTPFEFGFFRGLPE--NLFVLDIAGQIAFLLDIILHFFLAYRDPQTYRMVYKRTSIAIRYLKSSFLIDLLGCMPWDIIYKASGRKEEVRYLLWIRLYRVHKVTAFFRKMEKD  209
                     :*::***:***.**.**.**     *  ::. ** .*..*:.. * : ::*  :*:    .   **.:* ** *. *:. * *** :*     :* ** :* :** *  :     *:  :***
```

```
                                 S5                                          Pore-Helix                    S6
Arabidopsis_GORK      TRINYLFTRILKLLFVEVYCTHTAACIFYYLATTLPPENEGYTWIGSLKLGDYSYENFREIDLWKRYTTALYFAIVTMATVGYGDIHAVNLREMIFVMIYVSFDMVLGAYLIGNITALIV    311
Brassica_GORK         TRINYLFTRILKLIFVEVYCTHTAACIFYYLATTLPAENEGYTWIGSLKLGDYSYENFRKIDIWKRYTTSLYFAIVTMATVGYGDIHAVNLREMIFVMIYVSFDMVLGAYLIGNITALIV    310
Zingiber_GORK         IRINYLFTRIVKLIVVELYCTHTAACIFYYLATTLPASMEEYTWIGSLKLGDYSYSHFREIDLWRRYITSLYFAIVTMATVGYGDIHAVNTREMVFIMIYVSFDMILGAYLIGNMTALIV    335
Elaeis_GORK           IRINYLFTRIVKLIVVELYCTHTAACIFYYLATTLPASMEGYTWIGSLKLGDYSYSHFRDMDIAKRYITSLYFAIVTMATVGYGDIHAVNPREMVFIMIYVSFDMILGAYLIGNMTALIV    310
Phoenix_GORK          IRINYLFTRIVKLIVVELYCTHTAACIFYYLATTLPASMEGYTWMGSLKLGDYSYSHFRDIDLGKRYITSLYFAIVTMSTVGYGDIHAVNPREMVFIMIYVSFDMILGAYLIGNMTALIV    309
Oryza_GORK            IRINYLFTRIVKLIVVELYCTHTAACIFYYLATTLPESMEGYTWIGSLQLGDYSYSHFREIDLTKRYMTSLYFAIVTMATVGYGDIHAVNVREMIFIMIYVSFDMILGAYLIGNMTALIV    344
Brachypodium_GORK     IRVNYLFTRIVKLIVVELYCTHTAACIFYYLATTLPESMEGYTWIGSLKLGDYSYSNFREIDLAKRYMTSLYFAIVTMATVGYGDIHAVNVREMIFIMIYVSFDMILGAYLIGNMTALIV    326
Panicum_GORK          IRVNYLFTRIVKLIVVELYCTHTAACIFYYLATTLPESMEGYTWIGSLKLGDYSYANFREIDLAKRYITSLYFAIVTMATVGYGDIHAVNIREMIFVMIYVSFDMILGAYLIGNMTALIV    329
Papaver_GORK          IRINYLFTRIVKLLAVELYCTHTAACIFYYLATTLPAEKEGYTWIGSLKLGDYSYSNFREIDLGKRYIVSLYFAIVTMATVGYGEIHAVNVREMIFVMIYVSFDMILGAYLIGNMTALIV    331
Amborella_GORK        IRINYLFTRIVKLIVVELYCTHTAACIFYYLATTVPESEEGYTWIGSLTMGDYSYSHFREIDFIKRYLTSLYFAIVTMATVGYGDIHAVNPREMIFIMIFVSFDMILGAYLIGNMTALIV    314
Vulgaris_GORK         IRINYLFTRIVKLLTVELYCTHTAACIFYYLATTLPEREEGYTWIGSLTLGDYSYSHFREIDIWRRYTVSLYFAIVTMATVGYGDIHAVNMREMIFIMIYVSFDMVLGAYLIGNMTALIV    341
Chenopodium_GORK      IRINYLFTRIVKLLTVELYCTHTAACIFYYLATTIPEREEGYTWIGSLTLGDYSYSHFREIDLWRRYTTSLYFAIVTMATVGYGDIHAVNMREMIFIMIYVSFDMILGAYLIGNMTALIV    339
Lactuca_GORK          IRVNYLFSRIIKLIAVELYCTHTAACIFYYLATTLPAVEEGYTWIGSLKLGDYSYSNFREIDLWKRYTTSLYFAIVTMATVGYGEIHAVNMREMIFVMIYVSFDMVLGAYLIGNMTALIV    331
Daucus_GORK           IRIKYLFCRIVKLIVVEIYCTHTAACIFYYLATTLPAEKEGYTWIGSLKLGDYSYSNFREIDLWTRYITSLYFAIVTMVTVGYGDIHAVNLREMVFIMIFVSFDMVIGAYLIGNMTALIV    337
Medicago_GORK         IRVNYIIARIVKLLVVELYCTHTAACIFYYLATTLPESQEGYTWIGSLKLGDYAYSKFREIDIWKRYTTSLYFAIVTMATVGYGDIHAVNLREMIFIMIYVSFDMVLGAYLIGNMTALIV    336
Prosopis_GORK         IRINYMFTRIVKLIVVELYCTHSAACIFYYLATTLPPSQEGYTWIGSLQLGGYSYMNFREIDLWKRYTTSLYFAIVTMATVGYGDIHAVNMREMIFIMIYVSFDMVLGAYLIGNMTALIV    344
Nicotiana_GORK        IRINYLFTRIVKLITVELYCTHTAACIFYYLATTLSEQQEGYTWIGSLKLGDYSYSNFRDIDLWTRYTTSMYFAIVTMATVGYGDIHAVNLREMIFVMIYVSFDMILSAYLIGNMTALIV    319
Solanum_GORK          IRINYLFTRIVKLITVELYCTHTAACIFYFLATTLSEQQEGYTWIGSLKLGDYSYSNFRDIDLWTRYTTSMYFAIVTMATVGYGDIHAVNLREMIFVMIYVSFDMILSAYLIGNMTALIV    327
Coffea_GORK           IRINYLFTRIVKLIAVELYCTHTAACIFYYLATTLPPEKEGYTWIGSLKLGDYQYSSFRDIDLWKRYTTSMYFAIVTMATVGYGDIHAVNVREMIFIMIYVSFDMILGAYLIGNMTALIV    324
Sesamum_GORK          IRINYLFTRIVKLIAVELYCTHTAACIFYYLATTLPEEKEGYTWIGSLKLGDYSYAHFREIDIWKRYTTSMYFAIVTMATVGYGDIHAVNLREMIFIMIYVSFDMILGAYLIGNMTALIV    338
Cucumis_GORK          IRINYMFTRIVKLLVVELYCTHTAACIFYYLATTLPASEEGYTWIGSLKLGDYSYSHFREIDLWKRYTTSLYFAIVTMATVGYGDVHAVNLREMIFIMIYVSFDMVLGAYLIGNMTALIV    327
Vitis_GORK            TRINYMFTRILKLIAVELYCTHTAACVFYYLATTLPQSEEGYTWIGSLKLGDYSYSHFREIDIWKRYTTSLYFAIITMATVGYGDIHAVNLREMIFVMIYVSFDMILGAYLIGNMTALIV    318
Cannabis_GORK         IRINYIVTRIIKLIAVELYCTHTAACIFYYLATTLPPSKEGYTWIGSLKLGDYSYSSFRDIDLWKRYITSLYFAIVTMATVGYGDIHAVNMREMIFIMIYVSFDMILGAYLIGNMTALIV    338
Gossypium_GORK        IRINYLFTRIIKLIFVELYCTHTAACIFYYLATTLPREKEGYTWIGSLKLGDYSYSSFREIDLWKRYTTSMYFAIVTMATVGYGDIHAVNMREMIFIMIYVSFDMVLGAYLIGNMTALIV    329
                       *::*:. **:**: **:****:***:**:****:    * ***:*** :*.* *  **.:*:  **  .::****:** *****::**** ***:*:**:*****::.******:*****

                             C-Linker/A'       C-linker/B'     C-linker/C'    C-linker/D'        Helix A   β1     β2       β3
Arabidopsis_GORK      KGSNTERFRDKMNDLISFMNRKKLGRDLRSQITGHVRLQYDSHYTDTVMLQDIPASIRAKIAQLLYLPYIKKVPLFKGCSTEFINQIVIRLHEEYFLPGEVITEQGNVVDHLYFVCEGLL    431
Brassica_GORK         KGSNTERFRDKMNDLASFMNRKKLRGDIRSQITHHVRLQYDSKFTNTVMLQDIPASIRAKIAQLLYTPYIEKIPLFKGCSSEFINQIVVRLHEEYFFPGEVITEQGNVVDHLYFVCEGSL    430
Zingiber_GORK         KGSKTERFRDKMKDLLNYMNRNKLGKEIRDQIKGHVRLQYESNYTEASILQDIPISIRAKISQTLYRPYIERVPLFKGCSTEFVNQIVIRLQEEFFLPGEVLLEQGNAVDQLYFVCYGVL    455
Elaeis_GORK           KGSKTERFRDRMKELIKYMNRNKLGKDIRDQIKGHVRLQYESSYTEASVLQDIPVSIRAKISQTLYKPYIENVPLFKGCSAEFIHQIIIKLQEEFFLPGEVILEQGNAVDQLYFVCHGKL    430
Phoenix_GORK          KGSKTERFRDRMKDLVKYMNRNKLGKEIRDQIKSHVRLQYESSYTEASVLQDLPVSIRAKISETLYKPYIDNVPLFSGCSEEFIHQIIIKLQEEFFLPGEVLLEQGDAVDQLYFVCHGKL    429
Oryza_GORK            KGSRTERFRDKMKEVIRYMNRNKLGKDIREQIKGHLRLQYESSYTEASVLQDIPVSIRAKISQTLYKPYIESIPLFKGCSAEFIQQIVIRLQEEFFLPGEVILEQGSAVDQLYFVCHGAL    464
Brachypodium_GORK     KGSRTERFRDKMKEVIRYMNRNKLGKEIREQIKGHLRLQYESSYTEASVLQDIPISIRAKISQTLYKPYIESTPLFKGCSAEFIQQIVIRLQEEFFLPGEVILEQGSAVDQIYFVCHGEL    446
Panicum_GORK          KGSRTERFRDKMKEVIRYMNRNKLGKEIREQIKGHLRLQYESSYTEASVLQDIPISIRAKISQTLYKPYVESVPLFKGCSAEFIQQIVIRLQEEFFLPGEVILEQGSAVDQLYFVCYGAL    449
Papaver_GORK          KGSKTERFRDKMADIMKYMNRNKLGKEIRDQIKGHLRLQYESSYTEAVVLQDIPISIRAKISQTLYRDFIEDVPLFKGCSPELKNQIVIRLHEEFFLPGEVIMEQGNIVDQIYFVCDGVL    451
Amborella_GORK        KGSKTERFRDKMTDLIKYMNRNKLEKEIRDQIKGHVRLQYESSYTEASVLQDIPVSIRAEISQSLYKSTIEAVPLLKGCSSEFINQIVIRLHEEFFLPGEVILEQGNAVDQIYFISHGVV    434
Vulgaris_GORK         KGSKTERFRDKMTEVMKYMNRNRLSRELRSQIKGHLRLQYESSYTEASILKDIPISLRAKISQTLYTPYIGRGPLFKGCSSEFINQIVTRVHEEFFLPGEVILEQGNVVDQLYFVCHGLL    461
Chenopodium_GORK      KGSKTERFRDKMTEVMKYMNRNRLSRDLRNQIKGHLRLQYESSYTEASVLQDIPISLRAKISQSLYMPYIEKVPLFKGCSPEFINQIAIRVHEEFFLPGEVILEQGNVVDQLYFICHGLL    459
Lactuca_GORK          KGSKTERYRDRMTDLLKYMNRNRLGRDIRNQIKGHLRLQYESNYTDSTILRDLPISIRAKISQTLYKSYIEKVSLFKGCSLEFINQVVTKVHEEFFLPGEVIMEQGNAVEQLYFICHGEL    451
Daucus_GORK           KGSKTERYRDKMTDIIKYMNRNRLGRDIRNQIKGHLRLQYESTYTDAAALQDLPISIRAKISQTLYMSCVENVSLFKGCSSEFISQIVTRVHEEFFLPGELIMEQGNVVDQLYIVCHGLL    457
Medicago_GORK         KGSKTEKFRDRMTDLMKYMNRNKLHKDIREQIKGHVRLQFESSYTDAAVLQDIPISIRSKISESLYSPYIKNVSLFSGCSSEFINQIVTRLHEEFFLPGEVIMQQGYVVDQLYFVCDGVL    456
Prosopis_GORK         KGSKTEKFRDRMADLLKYMNRNKLGRDIREQIKGHMRLQYESSYTEPAAIQDLPISIRSKISQDLYLPYIEKVPLFKTCSSEFINQIVTRLHEEFFLPGEVILEQGNVVDQLYFVCHGVL    464
Nicotiana_GORK        KGSKTERYRDKMTDLLKYMNRNRLGRDIRNQIKGHLRLQYESAYTDAAVLQDIPISIRAKISQSLYQSYIENVPLFKGCSSEFISQVVTRVHEEFFLPGEVIMEQGNVVDQLYFVCHGVL    439
Solanum_GORK          KGSKTERYRDKMTDLLKYMNRNRLGRDIRSQIKGHLRLQYESAYTDAAVLQDIPISIRAKISQNLYQSYIENVPLFKGCSLEFISQVVTRVHEEFFLPGEVIMEQGNVVDQLYFVCHGVL    447
Coffea_GORK           KGSKTEKYRDKMTDLIKYMNRNRLGKDIRNQIRGHLRLQYESSYTDAAVLQDIPISIRAKISQSLYKSYIEEVILFKGCSAEFINQIVTRVHEEFFLPGEVIMEQGHAVDQLYFVCHGVL    444
Sesamum_GORK          KGSKTERYRDKMTDLIKYMNRNRLGKNLRNQIKGHLRLQYESSYTDASVLQDIPISIRAKISQTLYKSYVENVPLFNGCSSEFINQIVTRVHEEFFLPGEVIMEQGNVVDQLYFVCHGVL    458
Cucumis_GORK          KGSKTVKFRDKMADVMKYMNRNRLSREIRDQIKGHLRLQYESSYTEATVLQDIPISIRAKISQTLYLPYVQNVSLFRGCSPEFINQIVIRLHEEFFLPGEVIMEQGNVVDQLYFVCHGVL    447
Vitis_GORK            KGSKTERFRDKMTDVIKYMNRNRLDRDVRNQIKGHLRLQYESGYTEASVIQDLPISIRAKIAQTLYKPLVEKVSLFRGCSLELINQIVIRVHEEFFLPGEVIMEQGNVVDQLYFVCHGML    438
Cannabis_GORK         KGSKTEKFRDKMTDLIKYMNRNRLGKDIRNQIKGHVRLQYESSYTEASALQDIPISIRAKISQSLYLPFMEKVSLFRGCSSEFINQIVVKVHEEFFLPGEVIMEQGNVVDQLYFVCHGSL    458
Gossypium_GORK        KGSKTEKFRDKMADVIKYMNRNKLDRDLRNQIKGHLRLQYESTYTEAAVLQDIPISIRAKISQSLYLPYIENASLFKECSSEFINQIVIRLHEEFFLPGEVIMEQGNVVDQLYFVCHGVL    449
                      ***.* ::**:* ::  :***::*  ::*.**  *:***::* :*:   ::*:* *:*::*:: **   :    *:  ** *:  *:  :::**:*:***:: :**  *:::*::. * :
```

```
                           β4           β5    β6  Helix P     β7       β8     Helix B       Helix C                                         ANK1
Arabidopsis_GORK           EALVTKTDGSEESVTLLGPHTSFGDISIICNISQPFTVRVCELCHLLRLDKQSFSNILEIYFHDGRTILNNIMEEKE---SNDRIKKLESDIVIHIGKQEAELALKVNSAAFQGDFYQLK    548
Brassica_GORK              EALETKTDGTEDLVELLEPHTSFGDISIICNISQPFTIRVRSLCHLLRLDKQSFSNILEIYFHDGRKILNNLMEGKE---SNERIKKLESDIMIHIGKQEAELALKVNSAAFQGDIYQLK    547
Zingiber_GORK              EGITIGEDGSEETLLLLEPNSSFGEISILCNIPQPYAVRVRELCRLLRLDKQLFANIMEIYFVDGRTILSNLIEEN-E--SNGRIKHLESDITFHIGKQEAALALQVNSAAYYGDLLQLK    572
Elaeis_GORK                EGVGIGEDGP----LELEPNSSFGEIAILCNIPQPYTVRVCELCRLLRIDKQIFTNILQIYFVDGRKILSNLLEGN-D---SIQIKQLEGDIIFHIGKQEAELALRVNNAAFYGDLHYLK    542
Phoenix_GORK               EGVGIGEDGSKDTVLTLQPNSSFGEIAILCNIPQPFTVRVCELCRLLRIDKQVFTNILQIYFVDGRRILNNLLEGN-D---SNRIKQLESDITFHITKQEAELALRVNSDAFYGDLHHLK    545
Oryza_GORK                 EGVGIGEDGQEETILMLEPESSFGEIAVLCNIPQPFTVRVCELCRLLRLDKQSFTNILEIFFVDGRRILSNLSESS-E--YGSRIKQLESDITFHIGKQEAELTLRVNNAAFYGDMHQLK    581
Brachypodium_GORK          EGVGIGEDGQEETLLMLEPESSFGEIAILCNIPQPYSVRVCELCRLLRLDKQSFTNILEIYFVDGRKILSNLTENN-E--YGGRVKQLESDITFHIGKQEAELTLRVNSAAFYGDLHQLK    563
Panicum_GORK               EGVGIGEDGQEETLLMLEPESSFGEISILCNIPQPYTVRVCELCRLLRLDKQSFTNILEIYFVDGRRILSNLSE-S-E--YGGRVKQLESDITFHIGKQEAELILRVNSAAFYGDLHQLK    565
Papaver_GORK               EEMGIGEEGSEVTVSLLEPNSSFGELSILCNMPQPYTIRVCELCRLLRLDKESFISILEICSSDERIILTNLLEGKESNKDFHRMKQLESDVMFHIGKQEGELALRVNSAACHGDLYHLK    571
Amborella_GORK             EEVGIGEDGLEETIVQLEHDNSFGEIAILCNIPQPYTVRVCELCRLLRLDKQSFSNILQIYFIDARTVLNNLLEGK-E--SSLRLKQLESDIIFHINKQEAELALRVNSAAYYGDLYHLK    551
Vulgaris_GORK              EEVGIGQDGSEETVSRLEPGSTFGQISIFCNIPQPTTVRVLELCRLLRIDKESLSNIIDIYFYDGKKIFDNLLEGN-D--GKYNLKQVESDIASHIGKQESELALKVNNAAYHGDLYQIK    578
Chenopodium_GORK           EEVGIGQDGSEQIISRLEPNNTFGQISIFCNVPQPSIIRVIDLCRLLRIDKESLSNIIDIYFYDGKKIFDNLLEGNNN--GKYNLKQVESDIASHIAKQESELALKVNNAAHHGDLYQLK    577
Lactuca_GORK               EEIGICEDGSEETVSILKPHSSFGEVSILCNIPQPYTVRVCELCRLLRLDKQSFSNILEIYFHDGRKILNNLVEFN-A--SDARMKQLETDIILHIGKHEAELALRVNSSAFYGDLYQLK    568
Daucus_GORK                EEVGICEDGTEETVSVLRPNSLFGEVSILCNIPQPYTVKVSELCRLLRLDKQSFSNILEIYFHDGRKILNNLLEGK-E--SNLRLKQLESDITFHINKQEAELALRVNSAAYYGDLYQLK    574
Medicago_GORK              EEVGIADDGSEETVALLERNSSFGEISILCNIPQPHTVRVCELCRVLRIDKQSFSNILDIYFYDGKKVLDNLLEGK-E--S-IRGKQLESDITFHIGKLESELALKVNRTAFDGDLYQLK    572
Prosopis_GORK              EEVGPGENGSEETVSLLEPNSSFGQISILCNIPQPYTVRVCELCRLLRLDKQSFTDILDIYFYDGRKVLNNLLEGK-D--S-SRGKQLESDITFHIGKQEAELALKVNNAAFNGDLNHIK    580
Nicotiana_GORK             EEVGIAKDGSEETVALLEPNSSFGDISIVCNIPQPYTVRVCELCRLIRIDKQSFSNILEIYFHDGRRILSNLLQGK-E--SNLRVKQLESDIAFHIGKHEAELALKVNSAAYHGDLHQLK    556
Solanum_GORK               EEVGIAKDGLEETVSLLEPNSSFGDISIVCNIPQPYTVRVCELCRLLRIDKQSFANILEIYFHDGRRILSNLLQGK-E--SNLRVKQLESDIALHIGKHEAELALKVNSAAYHGDLHQLK    564
Coffea_GORK                EEVGTAKDGSEETLSLLEPNSVFGIVSILCNIPQPYTVRVCELCRLLRIDKQSFSNILEVYFHDGRTVLTNLLEEK-D--SAFSLKQLESDITLHIVKQEAELALKVNSAAYYGDLQQLK    561
Sesamum_GORK               EEVGIAADGSEDTVSLLEPNSLFGEISILCNIPQPYTVRVCELCRLLRIDKQSFSNILEIYFHDGRKVLTNLLEVI-T--D-----STISDITFHIGKQEAELALRVNSAAYYGDLYQLK    570
Cucumis_GORK               EELGIGQDSSEETIDLLQPNSSFGEISILCNIPQPYTVRVCELCRLLRIDKQSFTNILDIYFYDGRKILNNLLEGK-E--TNLRVKQLESDITFHIGKQEAELALKVNSAAYHGDLYQLK    564
Vitis_GORK                 EEIGIGADGSEETVLPLQPNSSFGEISILCNIPQPYTVRVLELCRLLRLDKQSFTDILEIYFYDGRRILNNLLEGK-E--SNLRVKQLESDITFHIGRQEAELALRVNSASYHGDLYQLK    555
Cannabis_GORK              EEVGIGEDGSEETVSLLQPSSSFGEISILCNIPQPYTVRVCELCRLLRLDKQSFTNILDIFFYDGRKILNNLLEVK-E--SNLRVKQLESDITFHIGKQEAELALKVNSSAFHGDLHQLK    575
Gossypium_GORK             EGIAIGEDGSEETVSLLEPNSSFGEISILCNIPQPYTVRVCELCRLLRLDKQSFSNILEIYFYDGRKVLNNLLEGK-E--SNLRVKQLESDISFHIGRQEAELALRVNCAAYNGDFYQLK    566
                           * :    :.       *   . ** :::.**: **  ::* .**:::*:**: : .*:::   * : :: *: :                *:  ** : *. * *:**  :  **:  :*

                                                   ANK2                            ANK3                          ANK4
Arabidopsis_GORK           SLIRSGADPNKTDYDGRSPLHLAACRGYEDITLFLIQEGVDVNLKDKFGHTPLFEAVKAGQEGVIGLLVKEGASFNLEDSGNFLCTTVAKGDSDFLKRLLSSGMNPNSEDYDHRTPLHVA    668
Brassica_GORK              SLVRSGADPNKTDYDGRAPLHLAASRGYEDITLFLIQEGVDINQKDKFGNTPLLEAVKAGQDRVIDLLVKEGASFDLEDAGNFLCTVVVKGDSDFLKRLLSSGMDPNTEDYDHRTPLHVA    667
Zingiber_GORK              GLVKAGADPKNTDYDGRSPLHLAASRGFEDITSFLIQECVDINVSDNFGNTPLFEAVKNGHDKVASLLYNNGAKLNLKDPSSHLCSAVARGDSDFIKRLLAYGIDLNLKDYDSRTPLHVA    692
Elaeis_GORK                GLIQAGADPNKTDYDGRSPLHLAAAKGHEDITLFLIQEGVNINLSDNFGNTPLFEAVKVGHDRVASLLFSRGAQLNLKDAGSHLCTAAAKGDSDFIRRALSYGVDPNSKDYDHRTALHIA    662
Phoenix_GORK               GLIQAGADPNKTDYDGRSPLHLAAARGHEDITLFLIQEGVDINLSDNFGNTPLFEAIKDGHDRVASLLFSRGAQLSLKDAGSQLCAAAAEGDSDFIKRALSYGVDPNSKDYDHRTALHIA    665
Oryza_GORK                 SLIRAGADPKNTDYDGRSPLHLAACKGFEDVVQFLLHEGVDIDLSDKFGNTPLLEAVKQGHDRVATLLFSKGAKLSLENAGSHLCTAVARGDTDFVRRALAYGGDPNARDYDHRAPLHIA    701
Brachypodium_GORK          GLIRAGADPKNTDYDGRSPLHLAASKGYEDVAQFLIHEGADINLIDKFGNTPLLEAVKQGHDRVATLLFKKGAILNLQNAGSHLCSAVSKGDSDFIRRALACGADPDSKDYDHRSPLHIA    683
Panicum_GORK               SLIRAGADPKNTDYDGRTPLHLAACRGYEDVVQFLLAEGVDINLTDQFGNTPLLEAVKQGHERVAALLFAKGAKLSLKNAGSHLCTAVAKGDSDFIRRALAFGADPNCGDYDHRTPLHIA    685
Papaver_GORK               RLIRAGADPNKTDYDGRTPLHIAASEGFEHIASFLIQEGVGVNIADKFGNTPLFEAVKRGNDRVASLLVKEGASLNIVDGGSCLCQAVSRGDSDFVKRLLSNGLDPNSKDYDQRTALHVS    691
Amborella_GORK             GLIRAGADPKKTDYDGRSALHLAASKGFEDITTFLIRETVDVNLTDKFGNTPLLEAIRYGHDRVASLLVKEGASLDIDDAGGFLCKAVASGNTDFLKRLLVYGMDPNSRDYDYRTALHKA    671
Vulgaris_GORK              NLLNAGADMNKTDYNGRSPLHIAAARGCEEIILLLVKEGVRINEKDNFGNTPLLEAVKNGNDKAAALLYREGARLDVKDVGSFLCTAVLGGDVDYLRRILANGIDPNSKDYDYRTPLHVA    698
Chenopodium_GORK           SLLNAGADLKKTDYNGRTPLHIAASGGCEDIVLFLIEEGADVNRTDNFGNTPLLEAVKNGHDKVAALLYKEKARIDVKDVGSFLCTVVLSGDVEFLKRILTNGMDPNSKDYDHRTPLHVA    697
Lactuca_GORK               SLIRAGADPNKKDYDGRTPLHLAASKGYEDITLFLIQEKVEVNVLDNFGNTPLLEAIKNGHDKVASILVKEGGSLKIDDAGWFLCTSVARGDIDYIRRILSNGVDPNSKDYDFRTPLHVA    688
Daucus_GORK                GLVRAGADPNKTDYDGRSPLHLAASKGYEDIILFLTQEGVDINISDNFGNTPLFEAIKSGHDQVVSLLIKEGAILKIGDSGPFLCTAVARGDYDLIKRVLSAGIDPNSSDYDHRTPLHVA    694
Medicago_GORK              SLIRAGADPKKTDYDGRSPLHLAACRGYEDITLLLIHERVDMNVKDNFGNTPLLEAVKNGHDRIASLLVREGASLKIDDGGSFLCSAVARGDSDYLKRLLSNGMDANLKDYDYRTPLHVA    692
Prosopis_GORK              GLIRAGADPNKADYDGRSALHLAASRGFEDITIFLIQEGVEIDLRDNFGNTPLLEAVKNGHDRVASLLVGKGASMKIEDAGSVLCNAVARGDSDYLKRLLSYGMDPNTKDYDYRSPLHVA    700
Nicotiana_GORK             SLIRAGADPNKKDYDGRSPLHLAASRGYEDITLFLIQEGVDINAPDKFGNTPLLEAIKSGHDRVASLLVKEGAFLKIENAGSFLCMVIAKGDSDLLRRLLSNGVDPNSKDYDQRTPLHVA    676
Solanum_GORK               SLIRAGADPKKKDYDGRSPLHLAASRGYEDITLFLIQEGIDINAPDKFGNTPLLEAIKIGHDRVASLLVKEGALLNIENAGSFLCMVIARGDSDLLRRLLSNGVDPNTKDYDQRTPLHVA    684
Coffea_GORK                NLIRAGADPNKNDYDGRSPLHLAASRGYEDIAQFLIQEGVDINVTDNFGNTPLLEAIKSGHEHVASLLVKEGASLQIENAGNFLCTVVAKGDSDLLRRFLTRGIDPNSKDYDQRTPLHVA    681
Sesamum_GORK               SLIRSGADPNKKDYDGRSALHLAASRGYEDVTLFLIQEGVDINAEDNFGNTPLLEAIKSGHDRVASLLSREGALLKIDNPGAYLCTVVSKGDTDLLRRVLANGIDPNSKDYDHRTPLHLA    690
Cucumis_GORK               GLVRAGADPNKTDYDGRSPLHLAASRGFEDIVVFLIQEGVNIDLKDNFGNTPLMEAIKNGNDQVAVLLSKEGASLKVDNPGSFLCTAVSRGDSDLVKRLLCYGIDPNSKDYDSRTPLHIA    684
Vitis_GORK                 SLIRAGADPNKTDYDGRSPLHLASARGFEDIVTFLIQEGVDVNISDNFGNTPLLEAIKNAHDRVASLLVNKGALLKIDDAGGFLCATIARGDSDFLKRILSNGIDPNSKDYDHRTPLHVA    675
Cannabis_GORK              SLIRAGADLNKTDYDGRSPLHLAASRGYEDITRFLIQQGVNVNAIDNFGNTPLLEALKNGHDRVSSLLVQEGASLMIDNAGSFLCTAVSRGDSDFLKRVLSNGVDPNSKDYDHRSPLHVA    695
Gossypium_GORK             SLIRAGADPNKTDYDGRSPLHLAASKGYEDITSFLIRHPVDINLKDKFGNTPLLEAIKNGHDNLAALLIKEGASLNIDDAGSYLCTAVAKGDSDFLRRLLSNGVDPNSRDYDHRTPLHVA    686
                            *:.:*** :: **:**: **:*:. * *.:  :*  .   ::  *:**:***:**:: .::    :*  . . : : : .  **     *: : ::* *  * : :  *** *: ** :
```

```
                     ANK5                              ANK6                                                                                    KHA

Arabidopsis_GORK     ASEGLFLMAKMLVEAGASVISKDRWGNSPLDEARLCGNKKLIKLLEDVKNAQSSIYPSSLRE-------LQ-EERIERRKCTVFPFHPQEAK----EERSRKHGVVWIPSNLEKLIVTA     776
Brassica_GORK        ASEGLFLMAKMLVEAGASVVAKDRWGNSPLDEARMCGNKKLIKLLEDADTSQPYIRPSSFHE-------PQ-DEKFERRKCTVFPFHPHE-------EPSRKHGVMVWLPRDLQKLVETA     772
Zingiber_GORK        ASVGFYAIAKMLSEAGASVFSVDRWGNTPLDEAMKCGSESMVMLLEDAKSQELSKFPERAQE-------LQ-EKMQSRRRCTVFPYHPRDL-----IESR-KEGVMLWIPHTLEELINSS     798
Elaeis_GORK          ASEGLYFIAKLLLEAGASVFAMDRWGATPLDEGRKSGNQSLMLLLEGAKSDELSKFPEHARE-------VQ-DKMHP-RRCTVFPFHPWDP-----KEAR-REGVVLWIPCTIEELIRSS     767
Phoenix_GORK         AAEGLYFIAKLLLDAGASVFAMDRWGATPLDEARKSGNKSLMLLLEGAKSDELSKFPERARE-------VQ-DKMHP-RRCTVFPFHPWDS-----KEER-REGVMLWIPCTMEKLIRSS     770
Oryza_GORK           AAEGLYLMAKLLVDAGASVFATDRWGTTPLDEGRRCGSRTMVQLLEAAKSGELSRYPERGEE-------VR-DKMHP-RRCSVFPHHPWDG-----GERR-REGVVVWIPHTIEGLVSSA     806
Brachypodium_GORK    AAEGLYMMAKLLVEAGASVFATDRWGTTPLDEGRKSGSKPLMMLLEQAKAEELSKFPARSEE-------VR-DKMHP-RRCSVFPNHPWDT-----DGKR-KEGVTLWIPHTIDWLIRSA     788
Panicum_GORK         AAEGLYLIAQMLVEAGASVFATDRWGTTPLDEARKCGGRTLLALLEQARAEELSKFPERGDE-------VR-DKMHP-RRCSVFPYHPWRAAAGTGAGRR-KEGVVLWIPHTIEGLVASA     795
Papaver_GORK         VSQGSYMIAKVLIEGGASVLTRDRWGNTPVDEGRTSGNKKLIKLLEEAESAELSTASTSGSAIDNTSELQVTNKANP-RKCTVFPFHPWDP-----KEKR-KEGVVLWVPPSVDELISMA     804
Amborella_GORK       AAEGLYSLAKNLLEAGASVFSKDRWGNTPLDEGHRSGSKLVINLLEEAKIDQLSRYPNYSQE-------IQ-GKIEA-KKCTVFPFHPWGP-----REER-KEGVVMWIPPTLDQLLSSS     776
Vulgaris_GORK        CSQGLYLMAKVLVEAGAYVTLKDRWGNTALDEAWMCGNKNLIKLLETAKSAQLSQSSGNIEE-------LS-DKKLQ-KKCTVFPFNPWGS-----KENR-RPGIVLWVPRTIDDLAKLA     803
Chenopodium_GORK     CSQGLYLMAKLLVEAGAYVTLKDRWGNTPLDEAWMCGNKNLIKLLEAAKAAQLAQSSDHIEE-------ML-DKKLH-KKCTVFPFHPWGT-----KENR-KPGIVLWLPRTIDELIRLA     802
Lactuca_GORK         ASQGSYIIVKLLVEAGASVLSKDRWGNTPLDEGRMSGNKMLIKLLEEAKKDVQLSELPEGSQE-------IT-DKMHP-RKCTVYAFHPWEE-----KDKI-TCGVVLWIPQTTEQLIKTA     793
Daucus_GORK          ASQGLYLLAKLLLESGASVISRDRWGNTPLDEGRMSGNKNLIKLLEEIAKSSQLAEISNQSPE-------IT-DKPQP-KKCRVYPFHPDKP-----KENKPSYGVMLWVPETMEELIKKA     800
Medicago_GORK        ASEGLIFMAKLLLEAGASVFTKDRWGNTPLDEARMSGNKNLIKLLEDAKSAQLTEFPF-PQE-------IT-DKVHP-KKCTVFPFHPWDP-----KEQR-RNGIVLWIPHTIQELIITA     796
Prosopis_GORK        AAEGLFLMAKLLIEAGASVFIQDRWGNTPLDEARMCGNKNLIELLEDARSAQLSRFPHLSSE-------LT-VKLHP-KKCTVFPFHPWGP-----KEHR-RHGIVLWVPHTIEELIETA     805
Nicotiana_GORK       ASQGLYSMAKLLLGAGASVFTKDRWGNTPVDEARVSGNKQMIGLLEEAKSAQLSEFPDVPHE-------IS-EKLRP-RKCTVFPFHPWES-----KDVR-KHGVVMWIPQTIEELVTTA     781
Solanum_GORK         ASQGQYSMAKLLLGAGASVFSKDRWGNTPVDEARVSGNKQMISLLEEAKSAQLCEFPDVPHE-------IS-DKLRP-RKCTVLPFHPWES-----KDLR-KHGVVLWIPQTIEELVTTA     789
Coffea_GORK          ASQGQYFMAKLLLDAGASVFARDRWGNTPLDEGRISGNKNMMDLLEEAESAQSSELSDLSQE-------STADKMLR-RKCTVFPFHPWEP-----KDAR-KCGVVLWVPDSIEELIREA     787
Sesamum_GORK         ASQGLYLMAKLLVEAGASVFSKDRWGNTPLDEGRLCGNKNMIRLLEEAKKAQLSEYPECSQE-------VT-DRKHT-KKCTVFPFHPWDP-----KEGR-KHGIVMWVPNTIDELIKAA     795
Cucumis_GORK         VSEGLTLMAKLLLESGASVFSKDRWGNTPLDEGRICGNKNMLKLLEEAKASQLSESPYSSRE-------FT-DKKPT-KKCTVFPFHPWDP-----EENK-RPGIMLWVPLTIEELIKES     789
Vitis_GORK           ASEGLYFMAKLLLEARASVFSKDRWGNTPLDEGWKCGNKNLMKLLEDAKVAQLSEFPDCSRE-------IT-DKMHP-RKCTVFPFHPWDP-----KEHK-RPGIMLWVPQTIEELIKTA     780
Cannabis_GORK        ASEGLYLMAKLLIEAGANVFSKDRWGNTPLDEGRMCGNKNLIKLLEDAKTAQLLDFPNSGRE-------IT-EKIHR-RKCTVFPFHPWDP-----KELR-RPGIVLWVPNSIEQLINTA     800
Gossypium_GORK       ASEGLYIMAKLLIEAGASVFSKDRWGNTPLDEARMCGNKNLIKLLEDAKSTQLSELPHCSKE-------FT-DKIHP-KKCTVFPFHPWDA-----KDQR-RHGIVLWVPHTMEALVTTA     791
                      : *   :.: *  . * *   **** : :**.  .*.. :: *** .   :                     .    ::* *   :*                *: :*:*   : *   :


Arabidopsis_GORK     AKELGLSDG-ASFVLLSEDQGRITDIDMISDGHKLYMISDTTDQT-------     820
Brassica_GORK        AQELGISNE-VPFVILSEEGGRITDIDMISDGQKLYLISDSTDQST------     817
Zingiber_GORK        QELLNCSVS----RLLSEEGGRILPAGLITDNLAPSVHAQNTTSTSDKELN-     845
Elaeis_GORK          REQLKSSGS----CILSEDGGRILDVDMISDGQKLYLVADQETTAHNNQHT-     814
Phoenix_GORK         REQLKSSGS----RILSEDGARILDVDMISDGQKLYLVADQETSTHNNQHT-     817
Oryza_GORK           QEKLGLAGSGEGLRLLGEDGARVLDVDMVHDGQKLYLVVGGGGDDGGTEARQ     858
Brachypodium_GORK    QEKLGLSGS--CLRLLGEDGARVQDVDMVNDGQKLYLVGDEDVGRSE-----     833
Panicum_GORK         QEKLGVPGPGARLRLLCEDGARVLDVDMVNDGQKLYLVGGDDDDQEDGE---     844
Papaver_GORK         SEHLQFYD--GS-RILSEDGGKILYIEMIHDGQKLHLV-------------     839
Amborella_GORK       SELLGCF---CT-CILSEDAGKILSVDMISDGQKLYAISDDSHIRDSL----     820
Vulgaris_GORK        SEKLNFPS--GS-CILSDDGGKILDIDMIENGQKLYLISETE----------     842
Chenopodium_GORK     SEKLNFLT--GS-CILSEDGGKILDVDMIENGQKLYLISETE----------     841
Lactuca_GORK         AEQLKLENYDTC-CIVTEDAGKILDVDMILDGQKLYLISEE-----------     833
Daucus_GORK          AEQLNCPQ--NS-CILSEDAAKILDADMISDGQKLYLIDETQ----------     839
Medicago_GORK        AEQIGFSS--DA-CILSEDAGKIIDISMIKDDQKLYLVNETH----------     835
Prosopis_GORK        AEQLNVSS--DC-SILSEDAGKITDANLIMDGQKLHLVHETH----------     844
Nicotiana_GORK       SEKLNFPS-D-S-CILSEDAGKILDIDMIADGQKLYLINDQST---------     821
Solanum_GORK         SEQLDFPS-GTS-CILSEDAGKILDVDMIVDGQKLYLINEST----------     829
Coffea_GORK          SKQLNFAS--CT-CILSEDAGKIIDVDMISNNQKLYLMNETD----------     826
Sesamum_GORK         SDQLESPN--GS-CILSEDGGKIFDVDMITDGQRLYMINETD----------     834
Cucumis_GORK         SEQLQVSG--EC-CILSEDGGKILDVHMIDESQKLYLVPDTH----------     828
Vitis_GORK           TEGLQFSS--ES-CILSEDGGKILDVDMISDGQKLYLLCETLDI--------     821
Cannabis_GORK        ADQLEISS--AA-CILSEDGGKILEEDMINDGQKLYLVGDTYSA--------     841
Gossypium_GORK       AEQLDLAG--AS-CMLTENGGKILDVDMINDGEKLYLIGGTH----------     830
                      . :          :: :: .::    :: :.
```

The cryo-EM structure of *At*GORK is used to restrict sequence gaps to inter-helical segments, with the superior coils and arrows defining extents of the secondary elements. The sequences include representative members from dicots and monocots: Arabidopsis thaliana (NP_198566.2), Brassica rapa (XP_009102317.1), Zingiber officinale (XP_042469382.1), Elaeis guineensis (XP_010905454.2), Phoenix dactylifera (XP_008795354.1), Oryza sativa Japonica Group (NP_001408448.1), Brachypodium distachyon (XP_003560852.1), Panicum virgatum (XP_039805885.1), Papaver somniferum (XP_026377173.1), Amborella trichopoda (XP_011629401.1), Beta vulgaris subsp. vulgaris (XP_010693307.2), Chenopodium quinoa (XP_021754949.1), Lactuca sativa (XP_023769988.1), Daucus carota subsp. sativus (XP_017218838.1), Medicago truncatula (XP_003616247.2), Prosopis alba (XP_028768743.1), Nicotiana tabacum (XP_016460239.1), Solanum lycopersicum (XP_004250206.1), Coffea arabica (XP_027077850.1), Sesamum indicum (XP_020551503.1), Cucumis sativus (XP_004140369.2), Vitis vinifera (XP_010660282.1), Cannabis sativa (XP_030505721.2), Gossypium hirsutum (XP_016749823.2).

**Figure S3. Sequence alignments for the representative GORKs from both dicots and monocots.**

### Figure S4

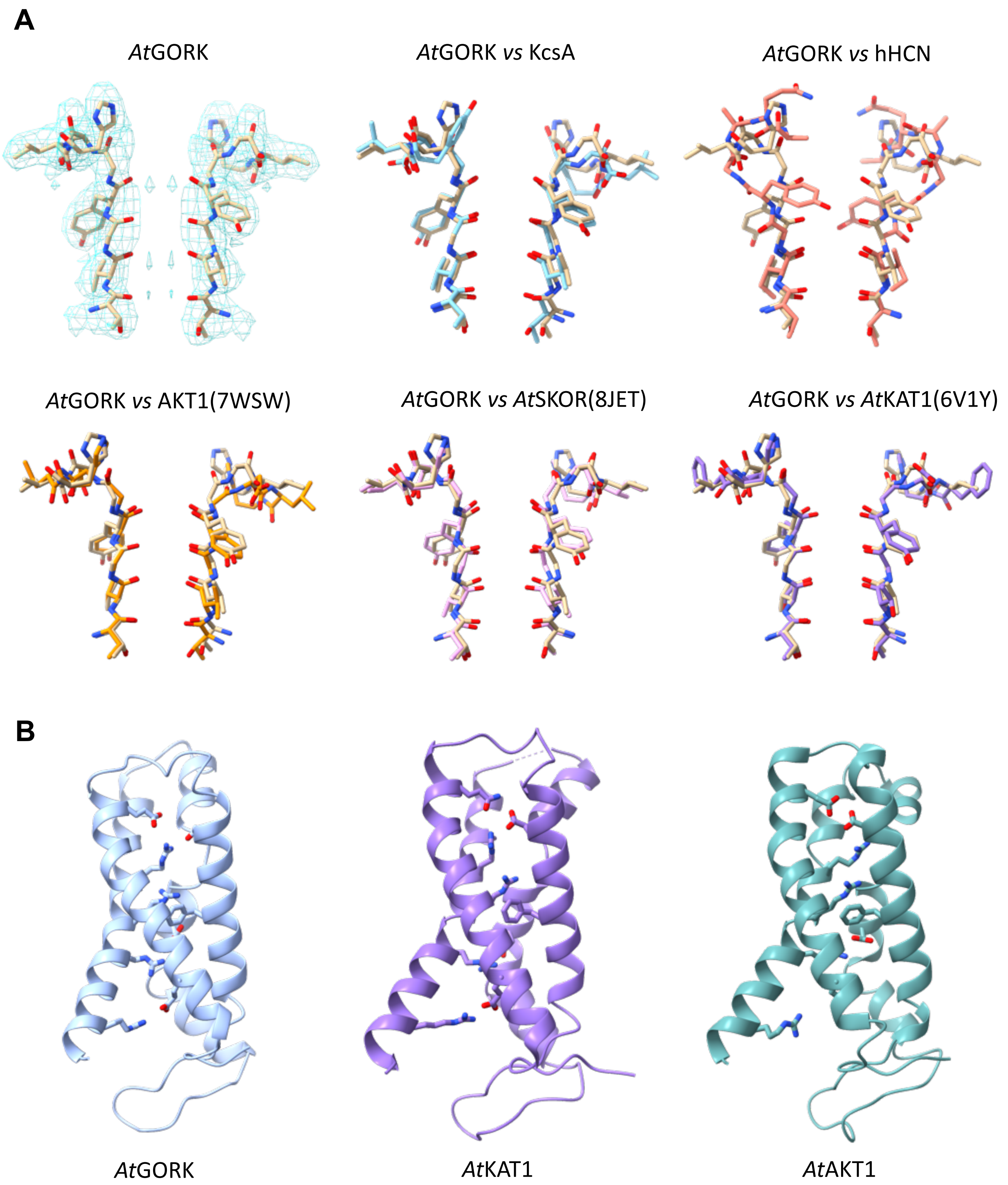


**Figure S4. Structural comparison of $K^+$-selectivity filter and VSD of *At*GORK with other $K^+$ channels.**

(**A**) Comparison of $K^+$-selectivity filters from *At*GORK with other $K^+$ channels. (**B**) Comparisons of voltage sensing domain (S1-S4) from *At*GORK (light-blue), AtKAT1(purple, 6v1y) and AtAKT1 (dark-green, 8wsw). The S4 helix in these three $K^+$ channel contains conserved positively charged residues, as shown in sticks, and adopt a resting "up" conformation.

### Figure S5

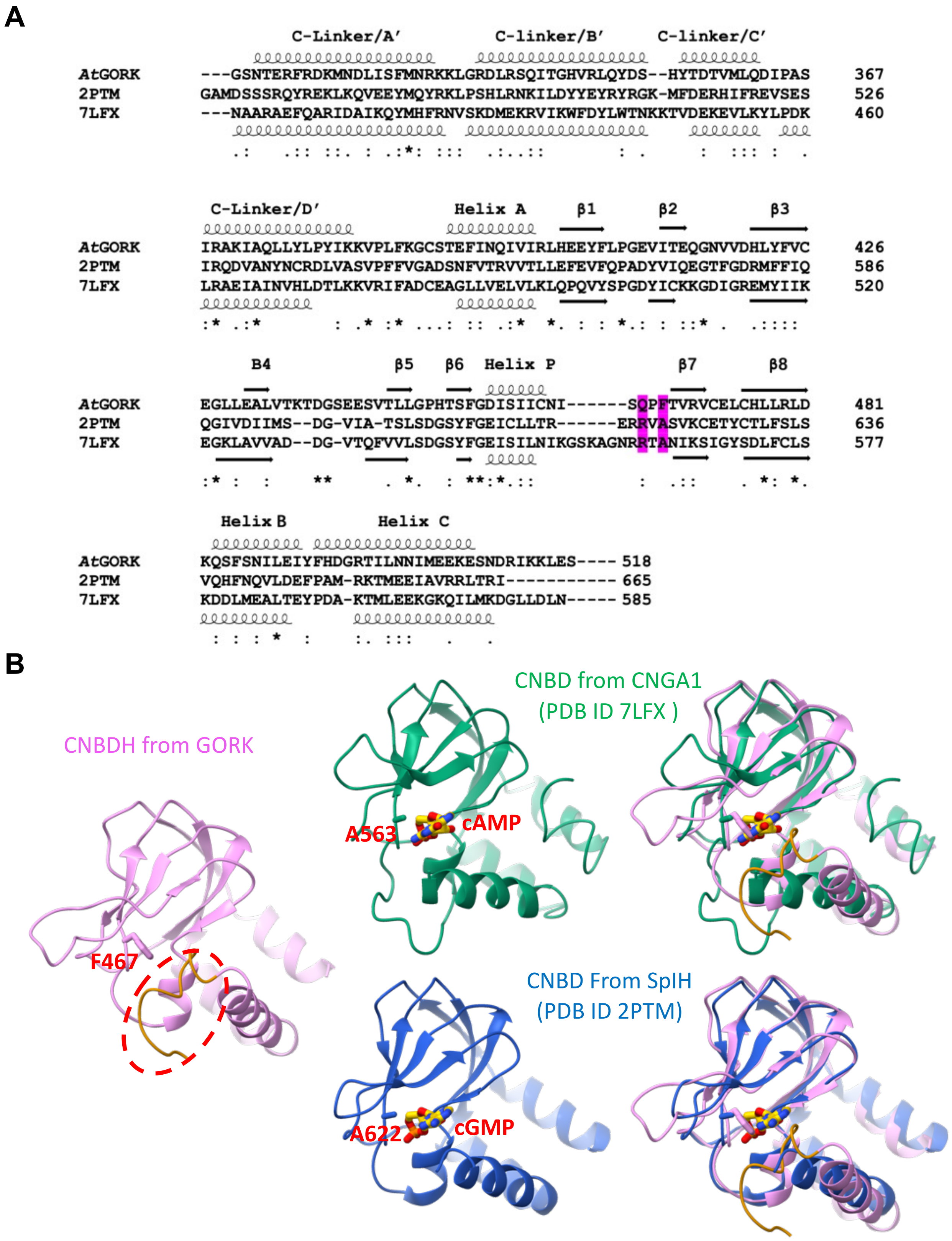


**Figure S5. Structural comparison of CNBD domain of *At*GORK with other K⁺ channels.**

(**A**) Structure-based sequence alignment of CNBDH of *At*GORK with CNBD from other $K^+$ channels. (**B**) Comparisons of CNBDH of *At*GORK with CNBDs from CNGA1 (7lfx, green) and SpIH (2ptm, blue). A small side-chain residue alanine (A563 in 7lfx, or A622 in 2ptm), located at the entrance of the cAMP/cGMP binding pocket, is replaced by bulky phenylalanine (F467 in *At*GORK), which likely blocks access for the secondary messenger molecules. Helix C adopts a closed conformation, with its adjacent connecting loop (red circle) to the next domain positioned at the entrance of the cAMP/cGMP binding site, indicating that large conformational changes would be required for cAMP/cGMP binding.

### Figure S6

**A**

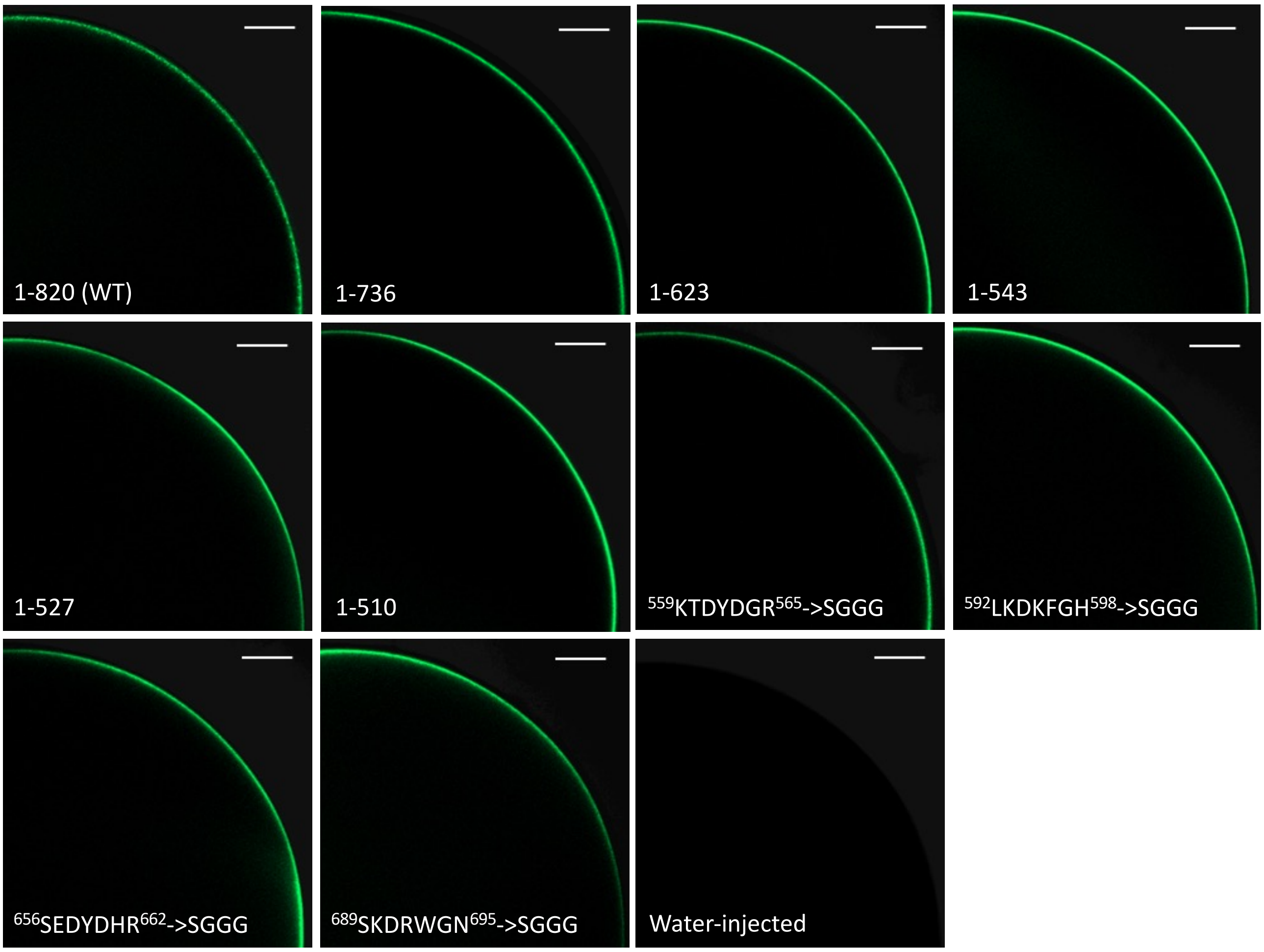


**B**

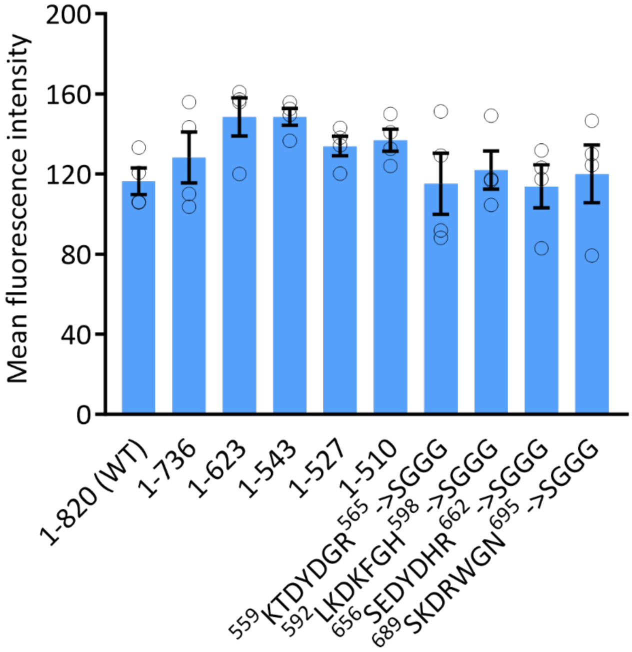


**Figure S6. Fluorescence and confocal imaging of the N-terminal GFP tagged *At*GORK.**

(**A**) Representative fluorescence images of *At*GORK wild-type (1-820) and mutants corresponding to Figure 3. The GFP fluorescence on the plasma membrane of oocytes was analyzed by LSM980 laser confocal microscope. The quarter of the whole cell is shown. Scale bars: 100 μm. (**B**) Mean fluorescence intensity measurements in ImageJ. A one-way ANOVA analysis was performed with a $P$ value of 0.1298. Data are mean ± SEM, $n = 4$.

### Figure S7

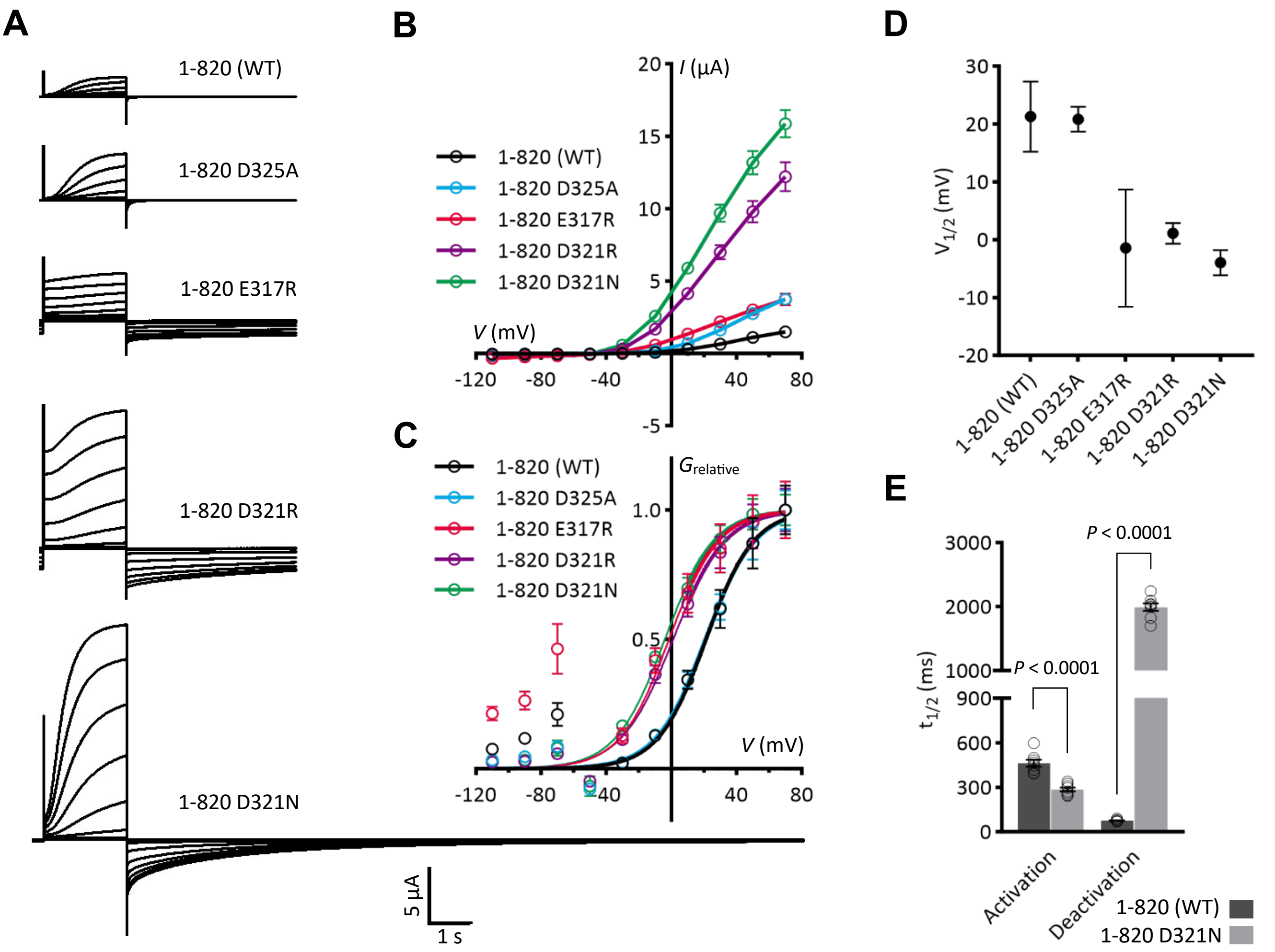


**Figure S7. Mutational tests of conserved acidic residues at the C-linker/TMD interface in full-length *At*GORK.**

(**A**-**E**) Electrophysiological analyses of E317, D321 and D325 mutations in *At*GORK$^{1\text{-}820}$. Representative current traces (**A**) and steady-state current-voltage (*I*-*V*) relations (**B**) are shown. Relative conductance-voltage ($G_{relative}$-$V$) curves (**C**) and half-activation voltage ($V_{1/2}$) values (**D**) were generated through Boltzmann sigmoidal fitting (outliers excluded). Conductance was calculated using the equation $G = I/(V - E_K)$, where $I$ is the steady-state current, $V$ is the test potential, and $E_K$ (–58 mV) was derived from the Nernst equation based on intracellular (~100 mM) and extracellular (10 mM) $K^+$ concentration in the oocyte TEVC recordings. Relative conductance was calculated by normalization to the maximal conductance. Halftime ($t_{1/2}$) for activation at +70 mV and deactivation at −110 mV, calculated as $\ln(2)\cdot\tau$ where $\tau$ is the time constant from single-exponential decay fitting, is shown in (**E**). Data are mean ± SEM, $n \geq 8$. Significance analysis was performed using unpaired Student's t-test, with *P*-values displayed on the bar charts.

### Figure S8

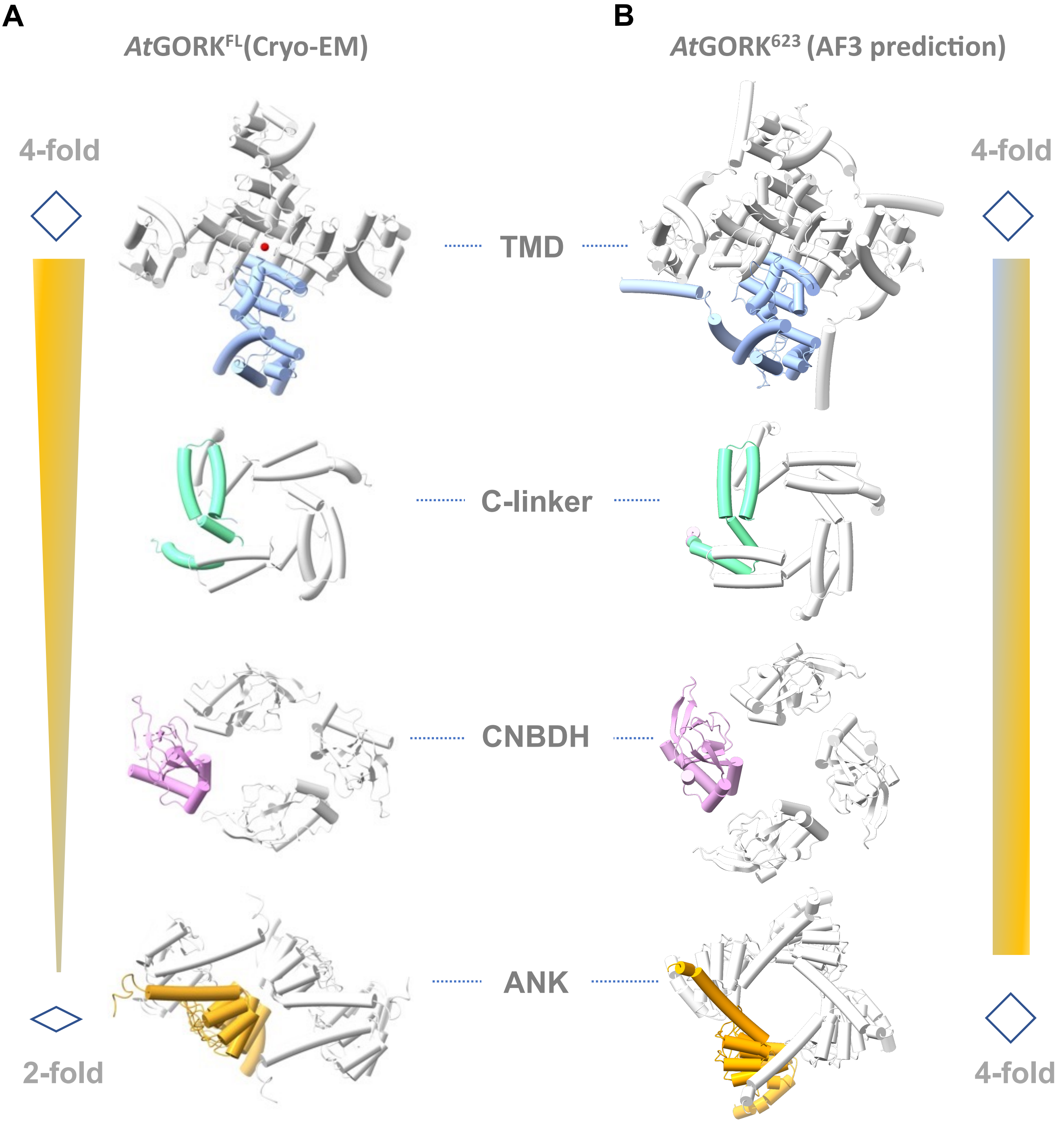


**Figure S8. Symmetry analysis of domain assembly in *At*GORK$^{FL}$(Cryo-EM) and *At*GORK$^{623}$(AF3 prediction).**

Table S1: Statistics of data collection, image processing, and model building

| **Sample** | $AtGORK^{FL1}$ | $AtGORK^{FL2}$ | $AtGORK^{623}$ | $AtGORK^{510}$ |
|---|---|---|---|---|
| **PDB** | 9KHF | 8WFZ | 9KHE | 9KHG |
| **EMDB** | 62338 | 37500 | 62337 | 62339 |
| **Data collection and processing** | | | | |
| Microscope | Titan Krios | Titan Krios | Titan Krios | Titan Krios |
| Detector | Gatan K3 | Gatan K3 | Gatan K2 | Gatan K3 |
| Magnification | 22,500 x | 22,500 x | 130,000 x | 22,500 x |
| Voltage (kV) | 300 | 300 | 300 | 300 |
| Electron exposure (e-/Å$^2$) | 50 | 50 | 50 | 50 |
| Nominal defocus range (µm) | -1.2 to -2.0 | -1.2to -2.0 | -1.2 to -2.0 | -1.2 to -2.0 |
| Frames per movie | 32 | 32 | 32 | 32 |
| Pixel size (Å) | 1.06 | 1.06 | 1.04 | 1.06 |
| **Reconstruction** | | | | |
| Software | cryoSPARC 3.2 | cryoSPARC 3.2 & Relion 3.0 | cryoSPARC 3.2 | cryoSPARC 3.2 |
| Symmetry imposed | C2 | C1 | C4 | C4 |
| Initial particle images (no.) | 18,650,563 | 18,650,563 | 1,139,470 | 2,840,921 |
| Final particle images (no.) | 156,313 | 39,551 | 75,626 | 141,755 |
| Map resolution (Å) | 3.4 | 4.3 | 3.2 | 3.3 |
| **Refinement and model validation** | | | | |
| CC (mask) | 0.7 | 0.76 | 0.82 | 0.84 |
| CC (box) | 0.63 | 0.66 | 0.58 | 0.64 |
| CC (peaks) | 0.55 | 0.45 | 0.54 | 0.62 |
| CC (volume) | 0.66 | 0.76 | 0.81 | 0.80 |
| RMSD Bond Length (Å) | 0.003 | 0.003 | 0.003 | 0.003 |
| RMSD Bond angles (degrees) | 0.632 | 0.701 | 0.627 | 0.572 |
| Favored (%) | 94.96 | 96.52 | 95.47 | 97.83 |
| Allowed (%) | 4.85 | 3.48 | 4.53 | 2.17 |
| Ramachandran plot outliers (%) | 0.19 | 0.00 | 0.00 | 0.00 |
| Molprobity score | 2.01 | 1.97 | 1.91 | 1.57 |
| Clash score | 14.33 | 17.44 | 11.95 | 10.17 |
